# Modality-specific and modality-general representations of subjective value in frontal cortex

**DOI:** 10.1101/2022.12.25.521898

**Authors:** Shilpa Dang, Jessica Emily Antono, Igor Kagan, Arezoo Pooresmaeili

## Abstract

Standard neuroeconomics theories state that the value of different classes of stimuli, for instance the hedonic value of food versus music, is transformed to a common reference scale that is independent of their sensory properties. However, adaptive behaviour in a multimodal dynamic environment requires that our brain also encodes information about the sensory features of rewarding stimuli. How these two processes, i.e. deriving a common reference scale for valuation and maintaining sensitivity to the sensory contexts for goal-directed behaviour, are implemented in the brain remains inadequately understood. Here, we acquired fMRI data while human participants were engaged in a dynamic foraging task requiring them to integrate the reward history of auditory or visual choice options and update the subjective value assigned to each sensory modality. Univariate fMRI analysis revealed modality-specific and modality-general value representations in orbitofrontal cortex (OFC) and ventromedial prefrontal cortex (vmPFC), respectively. Crucially, modality-specific value representations were absent when the task involved instruction-based rather than value-based choices. The analysis of effective connectivity further demonstrated that the modality-specific value representations emerged as a result of selective bidirectional interactions across the auditory and visual sensory cortices, the corresponding OFC clusters and the vmPFC. These results show how a valuation process that is sensitive to the sensory context of rewarding stimuli is enabled in the brain.

## Introduction

When we are presented with options for making a choice, our decisions are guided by our expectations of the rewards associated with each option. Theoretical frameworks of value-based decision-making suggest that our brain associates a subjective value to each available option based on their expected rewards (i.e., a valuation process), then compares these values, and makes a final choice (Balleine and Dickinson, 1998; Eryilmaz et al., 2017; Mannella et al., 2016; O’Doherty et al., 2017; Rangel et al., 2008; Valentin et al., 2007). In a multimodal dynamic environment, choice options can have fundamentally distinct sensory features and their associated values can change in time. For instance, the sound of a coffee machine, the smell of fresh bread, and the sight of a bottle of our favourite smoothie in the fridge could evoke the pleasant expectation of a nice breakfast and may therefore have similar value for us as we wake up in the morning. But after satiation we may not enjoy the smell of bread or the sight of smoothie as much as we may still be pleased by the sound of coffee machine.

To address the choice problem in a complex environment, as illustrated above, the brain’s valuation process should adhere to two fundamental principles. Firstly, it is important to be able to compare and choose between distinct types of outcomes, hence value representations independent of the specific type of rewards (e.g., juice, money, or social reward) and the stimuli predicting these rewards (e.g., sound of coffee machine or a picture of a smoothie bottle) should exist in the brain (Chib et al., 2009; Hampton et al., 2006; Hare et al., 2010; Levy and Glimcher, 2011; Lin et al., 2012; Noonan et al., 2011). This concept is often referred to as “common currency” coding of value (Berridge and Kringelbach, 2015; Levy and Glimcher, 2012; Montague and Berns, 2002; Padoa-Schioppa and Assad, 2008, 2006). Secondly, the encoding of recent stimulus-value associations should include specific information about the sensory features of each available option (Howard and Kahnt, 2021). These stimulus-specific representations of value enable flexible adaptation to rapid changes in internal states or external stimulus-value associations (Klein-Flügge et al., 2013; Nogueira et al., 2017; Rudebeck and Murray, 2014; Stalnaker et al., 2014; Wilson et al., 2014), which is crucial in a dynamic environment. Given that generalization across sensory features may conflict with the need for specific representations for each sensory context, the question arises as to how the brain reconciles these conflicting requirements while computing the subjective value. Here, we investigate this question in a dynamic choice environment in which the reward-predicting stimuli vary both in their sensory modality (auditory or visual) as well as their reward association history.

Orbitofrontal frontal cortex (OFC) and ventromedial prefrontal cortex (vmPFC) are key brain areas involved in the computation of subjective value and guidance of value-based choices (Montague and Berns, 2002; O’Doherty, 2014; Padoa-Schioppa, 2011; Rangel et al., 2008; Rolls, 2000; Setogawa et al., 2019; Stalnaker et al., 2015; Wallis, 2012). Different lines of evidence have pointed to the potential role of OFC, in particular the lateral OFC, in stimulus-specific valuation (Howard et al., 2015; Howard and Kahnt, 2017; Klein-Flügge et al., 2013; McNamee et al., 2013; Stalnaker et al., 2014), whereas vmPFC has been shown to underlie common currency coding of reward value (Chib et al., 2009; Hare et al., 2011, 2010, 2008; Lebreton et al., 2009; Levy and Glimcher, 2011; Lin et al., 2012; McNamee et al., 2013; Rolls et al., 2010; Smith et al., 2010). However, most of these insights have been derived from studies focused on visual stimuli and it remains uncertain whether they extend to the representation of the value of reward-predicting stimuli from different sensory modalities (but see Shuster and Levy, 2018). More importantly, the neural mechanisms that generate stimulus-specific representations of value and coordinate these with common currency valuation, have remained underexplored (Lim et al., 2013; O’Doherty et al., 2021). Stimuli from different sensory modalities have distinct representations at the input level – for instance in the early visual and auditory cortices. This feature provides a unique opportunity to selectively trace the neural mechanisms that generate the representations of value dependent or independent of the sensory context across the brain, from the early sensory areas to the frontal valuation regions.

In the current study, we aimed to find whether and how modality-specific stimulus value representations (SVR) exist in the key valuation regions when trial-by-trial updating of computed values of each sensory modality (auditory and visual) is necessary. Lateral and posterior regions of the orbitofrontal cortex receive highly specific and non-overlapping sensory afferent inputs from auditory and visual sensory areas (Barbas, 1993, 1988; Burks et al., 2018; Carmichael and Price, 1996; Martínez-Molina et al., 2019). More medial prefrontal areas including vmPFC on the other hand receive few direct sensory inputs (Carmichael and Price, 1996). Moreover, past research has shown that reward value modulates early sensory processing (Goltstein et al., 2013; Pleger et al., 2008; Rutkowski and Weinberger, 2005; Serences, 2008; Shuler and Bear, 2006). Based on these findings, we hypothesized that the representation of each option’s value should exist in OFC in a modality-specific manner and in vmPFC in a modality-general manner, and that the co-existence of these coding schemes in the frontal valuation regions is enabled through long-range interactions with the respective sensory cortices of each modality.

In order to test these hypotheses, we acquired fMRI data in a value-based decision-making task with a dynamic foraging paradigm adopted from a previous study (Serences, 2008). In this task, participants were presented with two stimulus options and chose the one they believed was associated with monetary value based on the trial-by-trial history of reward feedbacks. The two options were rewarded in an independent and randomized fashion to simulate foraging behaviour in a varying environment. To test the influence of sensory modality through which reward information was delivered, the task was performed under three different conditions: auditory, visual, and audio-visual, where the choice was made either intra-modally (between options from the same sensory domain) or inter-modally (between options form different sensory domains). To test the hypothesis that modality-specific representations in frontal areas were due to a difference in value processing requirements and not due to the difference in sensory processing requirements of the auditory and visual domains, a control task was also employed. The control task was designed in a way that the sensory processing requirements were identical to the value task, but selection was based on passively following an instruction as to which stimulus to choose, and not on the assessment of options’ reward history. Univariate fMRI analyses revealed modality-specific and modality-general value representations in lateral-posterior OFC and vmPFC, respectively. Effective connectivity analysis of a network consisting of regions exhibiting value modulations in auditory and visual sensory cortices, lateral and posterior OFC, and vmPFC, revealed how the two types of value representations emerge and guide value-based decisions.

## Results

### Behavioural results

We examined participants’ performance in two behavioural tasks (**Figure 1**), referred to as the value-based choice (value) and the instruction-based (control) tasks. In both tasks, participants aimed to maximize their performance while making a choice between two stimulus options. Specifically, in the value task, they sought to maximize their reward by choosing the stimulus option that they believed was associated with reward based on the feedback history (for details see Material and Methods). In the control task, participants’ objective was to enhance the accuracy of following instructions regarding which option should be chosen. The presented choice options were either from the same modality (two auditory stimuli or two visual stimuli or they were from different modalities (one auditory and one visual stimulus, as shown in **Figure 1A-B**). A choice was made either from stimulus set *S*_l_= {low pitch, green, auditory} or corresponding stimulus set *S*_2_ = {high pitch, red, visual}.

**Figure 1.**
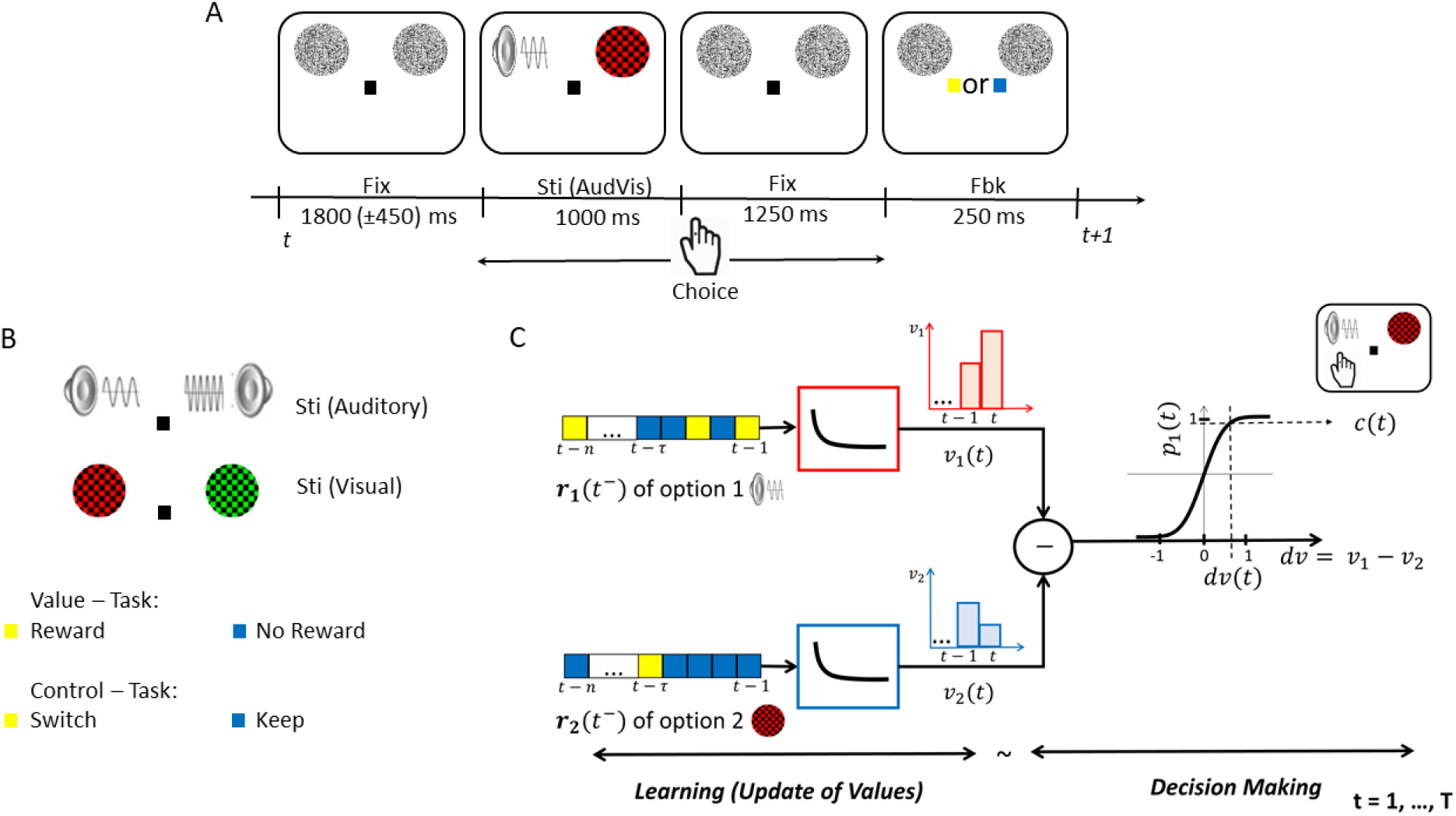
Experimental Paradigm and Computational Framework of Choice Behaviour. (**A**) General schematic of an audio-visual (AudVis) trial across both behavioural tasks (i.e., value and control tasks). After a jittered inter-trial interval of fixation (Fix), stimuli (Sti) options were presented. Participants made a choice during a response window of 2.25 s from the onset of stimuli, after which the fixation changed to either yellow or blue colour to indicate the feedback (Fbk). In all intervals, two white noise placeholders were shown on the screen. If a choice option was assigned to be from the visual modality, the corresponding placeholder was replaced with a coloured checkerboard (as shown here by a red checkerboard). (**B**) Stimuli options as presented during an auditory or a visual trial. In the value-based decision-making task, the yellow feedback indicated a reward and blue feedback no reward, whereas in the control task, the yellow and blue feedback instructed participants either to switch or to keep the past trial’s choice, respectively. (**C**) Reward history of option 1 and 2; i.e., *r*_l_(*t*^-^) and *r*_2_(*t*^-^); enter as inputs to two identical exponentially decaying filters that weigh rewards based on their time in the past and compute the subjective value of each option (i.e., *v_1_* and *v_2_*). The difference of the output of filters gives the differential value between the options (i.e., *dv*). The differential value determines the probability of choice between options (option 1 or 2, here option 1 is chosen as an example) according to a sigmoidal decision criterion.

In the value task, participants experienced an unpredictable outcome scenario with a dynamic reward structure (see Material and Methods). Reward baiting and change over delay (COD) strategies along with uninformed changes in the reward ratio across every block of trials motivated an exploratory choice pattern. Overall, the choice pattern in the value task exhibited matching behaviour nearly in accordance with the Herrnstein’s Matching Law (Herrnstein and Baum, 1970), which relates the choice behaviour to reward ratios {1:3, 1:1, 3:1}, as shown in **Figure 2A** for all modality domains (auditory, visual, audio-visual). Specifically, the choice ratios, which indicate the number of choices made towards one reward option (*S*_1_) over another (*S*_2_), increased as the *S*_1_: *S*_2_reward ratio increased. Importantly, the choice patterns were consistent across sensory modalities. This effect was captured by a strong main effect of reward ratio on choice ratio (*F*[2,38] = 183.8, p < 0.001) and no significant interaction between reward ratios and options’ sensory modality (*F*[4,171] = 0.95, p = 0.34) in a two-way repeated-measures ANOVA. Only a weak effect of modality on choice ratios was observed (*F*[2,38] = 5.95, p = 0.024), which corresponded to a tendency of participants to choose visual options more often than auditory options in the audio-visual block (for more details see the **Supplementary Information**). Therefore, we collapsed the choice ratios across modalities for a concise presentation of behavioural results (as shown in **Figure 2B**). Analysis of reaction times (RT) revealed no significant effect except for slower RTs in audio-visual compared to both auditory and visual conditions, reflecting that choosing between options from different sensory modalities is more difficult than choices made between items from the same modality (for details see the **Supplementary Information** and **nFigure S1**).

**Figure 2.**
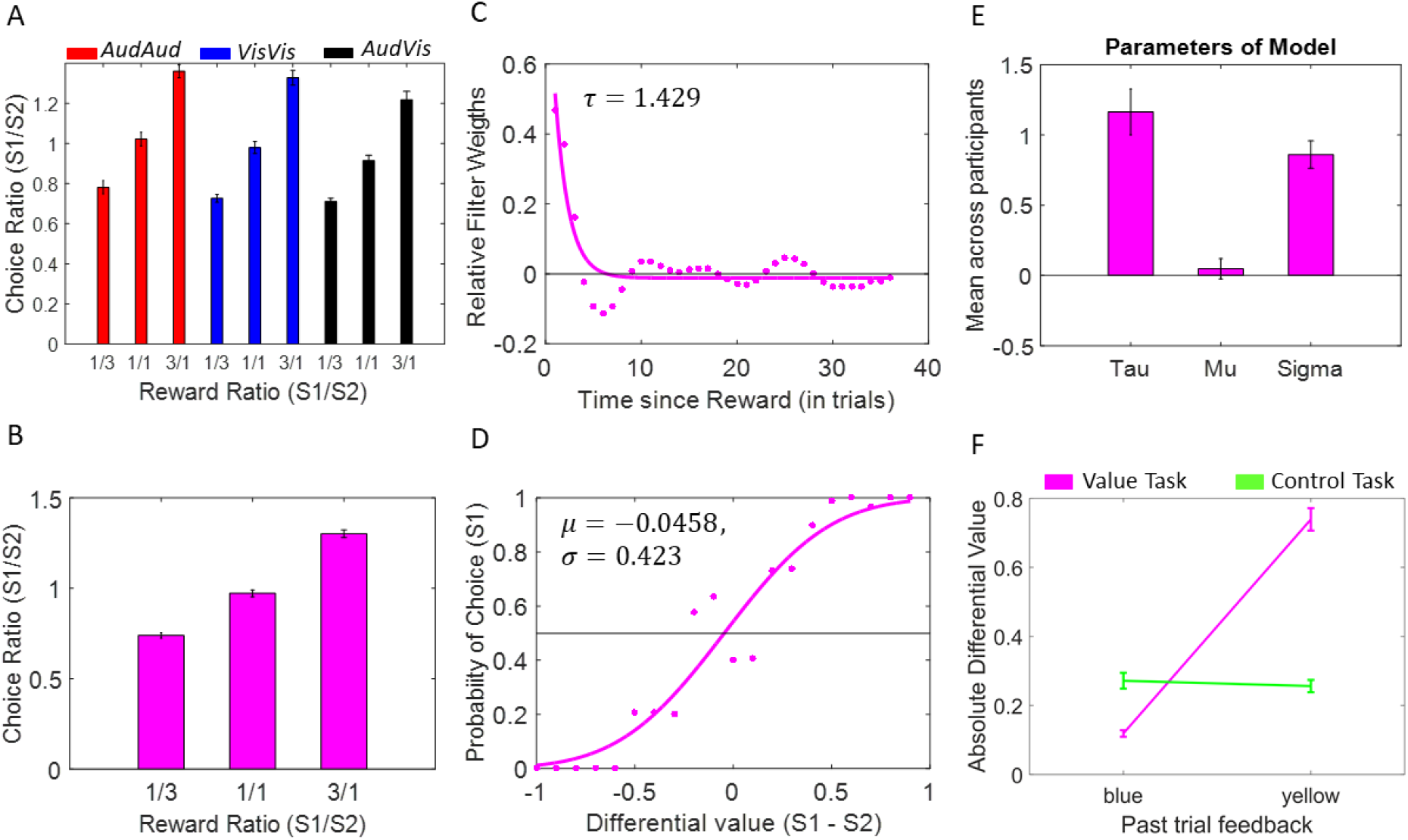
Behavioural Results. (**A**) Mean choice ratio across participants for each reward ratio {1:3, 1:1, 3:1} of options *S*_1_: *S*_2_ separately for each modality condition of the value task (*AudAud*, *VisVis* and *AudVis*). (**B**) Mean *S*_1_: *S*_2_ choice ratios in (A) collapsed across modalities for individual reward ratios. (**C**) Linear filter weights (dots) and exponential approximation (solid line) showing how past rewards are weighed based on their time in the past for a single participant in the value task. Parameter *τ* shows the timescale component of the best-fitting exponentially decaying function. (**D**) Mapping of differential value of option 1 and 2 to the probability of choice for option 1 (dots) and sigmoidal approximation (solid line) for the same participant as in (C). Parameters *µ* and *σ* of the best-fitting cumulative normal function show the participant’s biasness towards an option and sensitivity to value differences, respectively. (**E**) Mean parameters of the best-fitting curves across participants. (**F**) Relationship between feedback colours (yellow: reward / switch, blue – no reward / keep) and absolute differential value for the two tasks (value and control), across all participants. *S*_1_: *S*_2_ = {*low pitch:high pitch, green:red, auditory:visual*}, *S*_1_ - *S*_2_ = {*low pitch – high pitch, green – red, auditory – visual*}. Error bars indicate standard error of the mean (s.e.m.) across participants.

We next tested whether participants’ choices in the value task followed the predictions of our computational framework; i.e., they adhered to an LNP model (for details see Material and Methods). For this, we approximated the linear filter weights capturing the effect of reward history on choices (quantified by time scale parameter *τ*; **Figure 2C**), and we modeled the probability of choice function (quantified by biasness *µ*, and sensitivity to value differences *σ*; **Figure 2D**) for each participant. Across participants the mean time scale parameter *τ* was 1.22 (±0.15 s.e.m), which was significantly greater than zero *t*[19] = 8.39, p < 0.05, indicating that choices were in fact most impacted by recent rewards rather than distant rewards in the past (**Figure 2E**). Mean biasness *µ* across participants was 0.07 (±0.07 s.e.m), which was not significantly different than zero *t*[19] = 0.66, p = 0.52, indicating that participants did not have a bias towards any particular option. Finally, the mean sensitivity *σ* across participants was 0.81 (±0.10 s.e.m), which was significantly greater than zero *t*[19] = 8.81, p < 0.05 and insignificantly lesser than one *t*[19] = 1.95, p = 0.07, indicating that participants were aware of the value difference between options and had indeed adopted an optimal balance between exploration and exploitation (*σ* = 0 and *σ* ≫ 1 for extreme exploitative and explorative tendencies, respectively). Following this optimal strategy, participants were able to harvest 94.94% (±0.84% s.e.m) of the total rewards available. Overall, participants exhibited choice behaviours that were strongly predicted by the filter weights, estimated subjective values, and sigmoidal decision function of the LNP model in all conditions.

In order to demonstrate that the LNP model uniquely predicted the learning and choices in the value task, the fit parameters were also inspected for the data of the control task. Overall, in the control task participants passively followed the instruction provided by the feedbacks with a high accuracy (i.e., 95.2% ± 1.33% collapsed across keep/switch feedbacks), which indicated that they were aware of the task strategy. As choices in the control task were instructed, participants’ choices in this task were not expected to reflect tracking of option values beyond the instruction provided in the immediately preceding trial. This is exactly what we found when we compared participants’ beliefs about options’ value in the current trial depending on the type of feedback received in the past trial (blue or yellow), across the two tasks. The difference between the two tasks was captured by a significant interaction *F*[1,19] = 254.7, p < 0.001 between the task type (value or control) and feedback (yellow or blue) on determining the absolute differential values (*absDVs*; a measure of subjective preferences; **Figure 2F**). Indeed, *absDVs* showed a significant difference between the two types of feedbacks in the value task (mean±s.e.m. = 0.12±0.01 and 0.74±0.03 for blue and yellow feedbacks, respectively, p< 10^-ll^) but not in the control task (mean±s.e.m. = 0.27±0.02 and 0.26±0.02 for blue and yellow feedbacks, respectively, p = 0. 11). Analysis of the mean reaction times (RT) for the two types of feedbacks in the value and the control tasks revealed no significant mean or interaction effect (all p>0.05, for details see the **Supplementary Information**).

Collectively, our behavioural results confirmed that in the value task participants learned and updated their beliefs about options’ value from both sensory modalities through monitoring the feedbacks received on each trial, whereas in the control task they passively followed the instructions without any further processing of stimulus value, as intended.

### fMRI results

#### Modality-general and modality-specific stimulus value representations: vmPFC and OFC

We next specified a general linear model (GLM) for each participant to characterise the stimulus value representations (SVRs) of different experimental conditions in the brain and their potential dependence on the sensory modality of stimuli (see Material and Methods for the details of the GLM). Importantly, this GLM included several parametric regressors, referred to as the value regressors, that modelled the trial-by-trial modulations in participants’ beliefs regarding the value of each stimulus option as estimated from the computational modelling of the behavioural data (see the Material and Methods, **Figure 1C** and **Figure S3**). The contrasts that were defined based on these regressors hence assessed the extent to which fMRI responses changed as a function of modulations in subjective value of a stimulus. The stimulus value representations (SVRs) of each condition were then identified in the frontal cortex (see the Material and Methods and **Figure S2** for specifications of the search area) based on a group-level random-effects analysis on the contrast images obtained from all participants.

In intra-modal conditions (*AudAud* and *VisVis*), we examined the parametric value regressors separately for the auditory or visual sensory modality (referred to as *intraAudSV* > 0 and *intraVisSV* > 0 contrasts, respectively; for details see the Materials and Methods and **Table 1**). The auditory contrast in the value task revealed significant activations in vmPFC and left lateral OFC (latOFC) and the visual contrast revealed activations in vmPFC and left posterior OFC (postOFC, **Figure 3A-3E**). However, we did not find any significant activation in the right OFC for either of these contrasts. Lateralization of reward responsiveness in OFC could be related to a functional specialization of the left and right lateral OFC and has been reported in the past (Lopez-Persem et al., 2020). Importantly, we found a segregation of value-processing clusters across the sensory domains in OFC (*d* = 20.59 mm, *d*: Euclidean distance; criterion to decide a segregation was *d* > 8 mm according to the definitions in [54, 55]). On the contrary, the auditory and visual clusters in vmPFC were substantially overlapping (**Figure 3A, B, D**).

**Figure 3.**
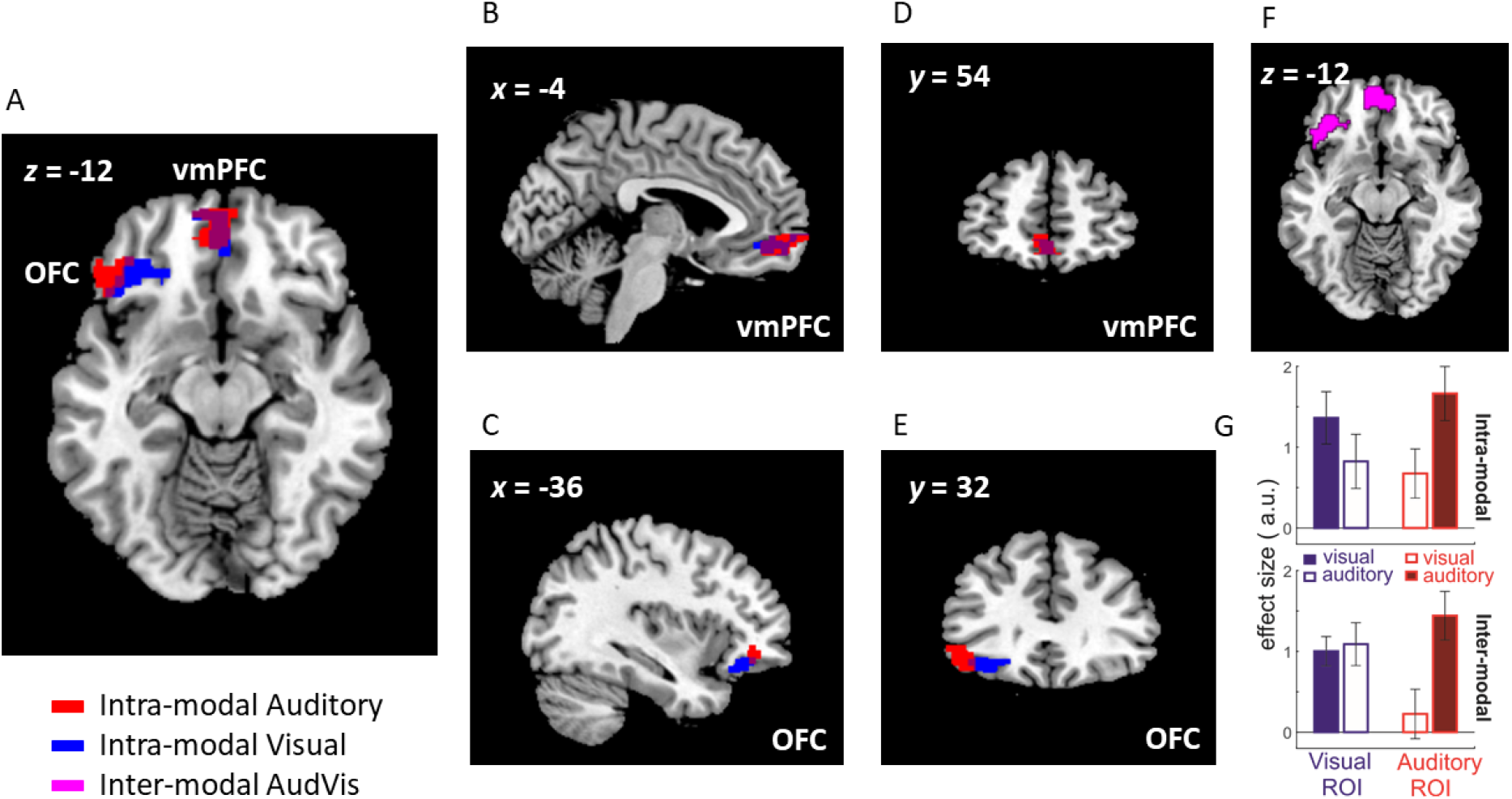
Stimulus value representations (SVR) across different sensory modalities. (**A-E**) In intra-modal conditions, segregated modality-specific SVRs in OFC for auditory modality in left lateral OFC (red cluster) and for visual modality in left posterior OFC (blue cluster); and overlapping modality-general SVRs in vmPFC for auditory and visual modalities, were found. (**F**) In inter-modal condition, SVRs were found in OFC and vmPFC. All cluster activations shown here are significant at SVFWE corrected P < 0.005. (**G**) To test the robustness of the modality-specificity of SVRs in the lateral OFC, visual and auditory ROIs were defined based on the data of intra-modal condition and tested on the data of inter-modal condition that was recorded in separate runs. Each bar represents the effect size (t-value) of the parametric value regressors. Blue and red bars depict the effect size in the visual and auditory ROIs (extracted from the blue and red clusters in Figure 3A-E). Filled bars show the responses to the stimulus value from the same modality as the ROI and unfilled bars show the responses to the stimulus value in a different modality.

**Table 1.** Stimulus value representations in vmPFC and OFC for various contrasts.

|  |  |  |  |  |  |  |
| --- | --- | --- | --- | --- | --- | --- |
| Contrast | Region | X | Y | Z | *t*(19) | *k* |
| *intraAudSV* > 0 | vmPFC_L | -6 | 64 | -2 | 5.33 | 121 |
|  | latOFC_L | -48 | 32 | -10 | 5.31 | 134 |
| *intraVisSV* > 0 | vmPFC_L | -8 | 58 | -14 | 4.70 | 142 |
|  | postOFC_L | -30 | 26 | -18 | 6.20 | 143 |
| *interAudVisSV* > 0 | vmPFC | 0 | 52 | -10 | 6.93 | 537 |
|  | antOFC_L | -36 | 36 | -14 | 7.60 | 242 |

MNI coordinates (x, y, z) and T value corresponds to the local maxima peak of the cluster activations at SVFWE corrected $P < 0.005$ (cluster labels are from AAL atlas (Rolls et al., 2020)). Statistical maps were assessed for cluster-wise significance using a cluster-defining threshold of $t(19) = 3.58$ , $P = 0.001$ ; and using small volume corrected threshold of $P < 0.005$ (referred to as a small volume family-wise-error (SVFWE) correction) within the frontal search volume.

The aforementioned results hinted towards a functional segregation in the OFC, but not vmPFC, with regard to the representations of stimulus value from the visual and auditory sensory modalities. If this is the case, we should observe both types of representations (i.e., SVRs for visual and auditory modality) in the inter-modal condition where the values of both sensory modalities are simultaneously tracked. This is exactly what we observed when we examined the modulation of fMRI responses by the value modulations of the inter-modal condition (*AudVis*). This analysis (referred to as *interAudVisSV* > 0 contrast in **Table 1**) showed significantly activated clusters in vmPFC and in the left OFC (**Figure 3F**). Interestingly, in the activated cluster in the left OFC, there were two local maxima peaks (-44, 26, -12, t(19) = 5.24 and -36, 24, -18, *t*(19) = 4.61), which were adjacent (d < 8mm) to the respective auditory and visual peaks found in the intra-modal conditions (see **Table 1**).

To test whether the observed segregation of auditory and visual SVRs in the lateral OFC in intra-modal conditions truly reflected a functional specialization for modality-specific valuation, we next performed a cross-validation procedure. To this end, we defined ROIs consisting of the most modality-specific voxels of the respective SVRs in the intra-modal conditions (red and blue OFC clusters shown in **Figure 3A**, see Material and Methods for a detailed description of voxel selection) and tested the extent to which modality-specificity generalized to the inter-modal condition (**Figure 3G**). As expected, we observed a strong interaction effect (*F*[1,19] = 23.32, p < 0.001) between the ROIs (visual or auditory) and the sensory modality of options (same or different modality) in intra-modal condition (**Figure 3G**, upper panel), reflecting that each ROI contained voxels that selectively represented the value of stimuli from the same modality. Importantly, this selectivity also generalized to the inter-modal condition, as we observed a significant ROI x Modality interaction (*F*[1,19] = 4.86, p = 0.04, see **Figure 3G**, lower panel) in this condition as well. Repeating the same analysis in the vmPFC revealed no generalization of ROI x Modality interaction to the inter-modal condition (*F*[1,19] = 0.22, p = 0.89). Since intra- and inter-modal conditions were recorded in different fMRI data acquisition runs, this result provides evidence that the clustering of modality-related value representations in the lateral OFC reflected a genuine specialization of the neural responses. However, only the auditory cluster in the OFC seemed to preserve its modality-specificity of responses across runs (p = 0.025 for the post-hoc pairwise comparison between responses to the same versus different modality). This pattern suggests that the auditory stimuli not only activated the auditory-specific SVRs but also to some extent coactivated the visual SVRs. This is likely because the visual placeholders which were presented on the screen in all intervals during a trial were utilized by the participants as spatial landmarks to facilitate the localization of the auditory tones to the left or the right side (see **Figure 1A**).

An alternative explanation for the segregation in modality-wise representations in the OFC is that rather than reflecting the functional specialization of neural responses for the visual and auditory values, they reflected differences in the sensory properties of stimuli. To rule out this possibility, we next examined the control task (see **Table S1**, **Table S3** and **Figure S4** in the **Supplementary Information**). Crucially, the modality-specific activations in OFC were absent in the control task when the same contrasts as in the value task were examined, demonstrating that OFC representations reflect the trial-by-trial updating of stimulus-value associations rather than the sensory features of stimuli or choice based on the instruction (**Table S3**). On the contrary, in the control task activations overlapping with the modality-general representations in vmPFC were found (**Figure S4**). Inspecting contrasts which explicitly tested the value and the control task against each other supported these results (**Table 2**). Contrasting the unmodulated trial identity regressors which modelled the responses to different sensory modalities revealed no significant difference between the two tasks, confirming that the control task was powerful enough to capture the sensory responses to the stimuli. Contrasting the parametric regressors however showed strong bilateral activations in postOFC in the value task, supporting the involvement of the OFC in representing the value of stimuli beyond their sensory properties. The lack of activations in the vmPFC for this contrast further highlights a general role of this area in representing the final choice irrespective of whether or not choices were informed by value or were instructed.

**Table 2.** Stimulus value representations in OFC for various interaction contrasts.

| Contrast | Region | *X* | *Y* | *Z* | *t*(19) | *k* | *SVFWE*<br>*corr* |
| --- | --- | --- | --- | --- | --- | --- | --- |
| *Value_all* *Parametric* > *Control_all* *Parametric* | postOFC_L | -24 | 24 | -12 | 6.29 | 584 | **P < 0.001** |
|  | postOFC_R | 30 | 24 | -10 | 7.42 | 524 | **P = 0.001** |
| *Value_all* *Trial identity* > *Control_all* *Trial identity* | No surviving clusters |  |  |  |  |  |  |

Statistical maps were assessed for cluster-wise significance using a cluster-defining threshold of $t(19) = 2.86$ , $P = 0.005$ ; and using small volume corrected threshold of $P < 0.005$ (referred to as a small volume family-wise-error (SVFWE) correction) within the frontal search volume. MNI coordinates (*x*, *y*, *z*) and *T* value corresponds to the local maxima peak of the cluster activations and activations shown in bold survive a SVFWE corrected $P < 0.005$ .

Together, the findings of our univariate analyses provide evidence that the valuation of stimuli from auditory and visual sensory modalities is confined to segregated loci in the OFC. However, the pattern we observed is suggestive of a partial modality-specificity in our task, as we found evidence for some degrees of coactivation between the visual and auditory clusters. Additionally, our results indicate that the representation of stimulus value is independent of sensory modality in the vmPFC and that this region is involved in processing information related to the final choice across different tasks.

#### Stimulus value representations outside of frontal valuation regions: Whole-brain analysis

In order to identify regions exhibiting value modulations outside the valuation regions, specifically in auditory and visual sensory cortices, we performed a whole-brain analysis using the GLM described previously. For this purpose, we estimated the overall effect of the parametric value regressors across all conditions in the value task (*AudAud*, *VisVis* and *AudVis*), which revealed bilateral activations in the auditory and visual cortices (whole-brain FWE corrected P < 0.05, cluster size *k* > 10 voxels; **Table 3**, **Figure S5A** and **S5B**). When estimating the value modulations for individual conditions separately (auditory, visual), we found modality-specific activations in respective sensory cortices only (see **Figure S5C-D**), whereas in intermodal condition both sensory cortices were activated (see **Figure S5E-F**).

**Table 3.** Stimulus value representations outside of valuation regions.

| Table 3. Stimulus value representations outside of valuation regions |  |  |  |  |  |
| --- | --- | --- | --- | --- | --- |
| Region | *X* | *Y* | *Z* | *t*(19) | *K* |
| SFGmed | 0 | 54 | 40 | 10.50 | 64 |
| Caudate_L | -14 | 24 | 8 | 10.37 | 67 |
| **visSen_L** | **-20** | **-90** | **2** | **9.75** | **81** |
| **visSen_R** | **24** | **-96** | **10** | **8.86** | **119** |
| Hippoc_L | -36 | -38 | -12 | 9.62 | 30 |
| Hippoc_R | 34 | -22 | -12 | 9.51 | 29 |
| **audSen_L** | **-66** | **-30** | **-4** | **9.08** | **24** |
| **audSen_R** | **66** | **-12** | **-4** | **8.99** | **15** |
| Angular_L | -42 | -56 | 26 | 8.62 | 55 |
| MNI coordinates (x, y, z) and T value corresponds to the local maxima peak of the cluster activations at FWE corrected $P < 0.05$ (cluster labels are from AAL atlas (Rolls et al., 2020)). SFGmed – medial Superior Frontal Gyrus; Hippoc – Hippocampus; visSen – Visual Sensory Cortex; audSen – Auditory Sensory Cortex. Highlighted (in bold) activations were used as ROIs in the effective connectivity analysis. | | | | | |

In addition to sensory cortices, we found significant value modulations in areas involved in processing different aspects of value-related information, such as detecting the reward prediction errors (Caudate Nucleus), formation of memories about past events (Hippocampus), selection of action sets (SFGmed/dmPFC) (Rushworth et al., 2004) and processing of symbolic information related to monetary value (Angular gyrus). Since the specific aim of the current study was to shed light on how modality-specific and modality-general valuation is coordinated across the frontal and sensory areas, we only included the whole-brain activations that were located in early visual or auditory areas in our subsequent effective connectivity analyses.

#### Modality-specific effective connectivity between sensory and valuation areas

We next examined the effective connectivity (EC) of a network consisting of sensory and valuation regions that showed significant value-related modulations in our univariate analysis. This network comprised 5 ROIs as shown in **Figure 4** (i.e., auditory and visual sensory areas: *audSen* and *visSen*; distinct modality-specific auditory and visual SVRs in the OFC: *audOFC* and *visOFC;* and overlapping modality-general SVRs in the *vmPFC*, see Material and Methods for details). The EC analysis (Friston et al., 2003) provides an estimation of the degree to which different connectivity patterns across this network contribute to the generation of value representations. Importantly, the EC analysis allows for an additional test of our main hypothesis regarding the existence of modality-specific representations of stimulus value across the brain, as models with and without a specific connectivity pattern for auditory and visual values were tested against each other (see the model space in **Figure 4A**). We then looked for the most probable connectivity pattern in this network using a Bayesian model comparison approach (Stephan et al., 2009).

**Figure 4.**
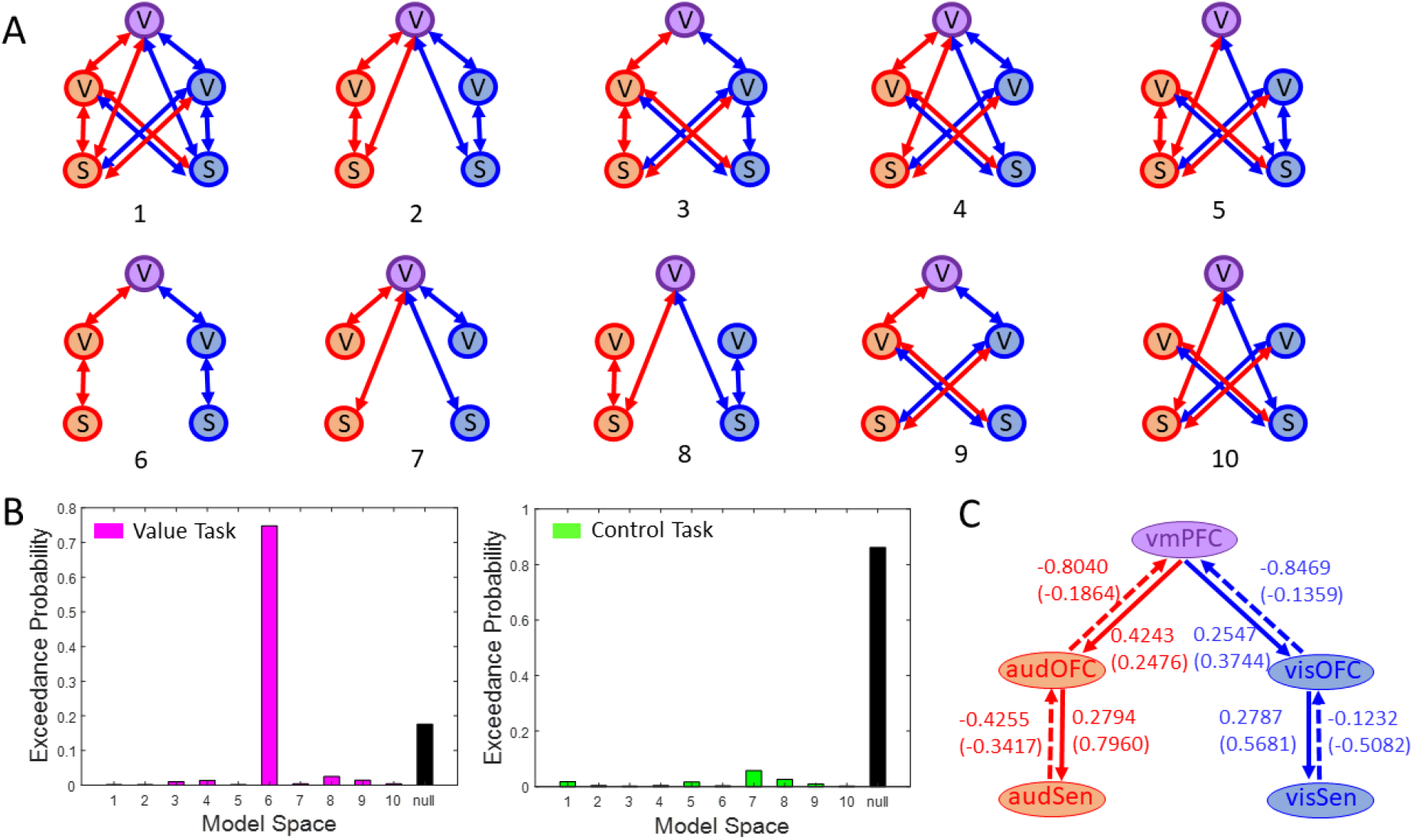
Modality-specific effective connectivity. (**A**) Model space consisted of 10 biologically plausible models per task (value or control). Models differed in their modulatory connections between nodes of a network comprising 5 ROIs (ROIs: V – Valuation region, S – Sensory region, purple V – vmPFC, red V – Auditory OFC, blue V – Visual OFC, red S – Auditory Sensory Cortex, blue S – Visual Sensory Cortex, see also panel C). Modulatory connections in red exist during conditions in which an auditory stimulus was selected and those in blue exist during conditions in which a visual stimulus was selected in both intra- and inter-modal conditions. (**B**) Exceedance probabilities for all connectivity models in the value and control task. In the value task, Model 6 is the most likely model, with exceedance probability of 0.75. In the control task, the null model, which did not have modulatory connectivity between ROI pairs for any condition was the winning model. (**C**) The weights of feedforward (dashed) and feedback (solid) modulatory connections in the winning model of the value task. Connection weights are shown for conditions in which an auditory (red) or a visual (blue) stimulus either in intra-modal or inter-modal conditions (in brackets) was selected. All parameters were significant at posterior probability of P > 0.99.

In the value task, our findings revealed that the most probable model was one that included modulatory connections between the sensory cortices, modality-specific clusters in the OFC, and vmPFC (i.e., model 6, **Figure 4A-B**). The winning model in the value task contained two distinct valuation sub-networks: an auditory sub-network comprising audSen, audOFC and vmPFC, and a visual sub-network with visSen, visOFC and vmPFC as nodes (**Figure 4C**). Moreover, the sensory cortices did not directly communicate with vmPFC (models 2, 4, 5, 7, 8 and 10) and we did not find evidence for the cross-modality of the connectivity between the sensory cortices and the value regions in OFC (models 1, 3, 4, 5, 9 and 10, containing connections between visual cortex and auditory OFC and auditory cortex and visual OFC). Additionally, in the control task, the null model, which lacked any modulatory connections between ROIs, yielded the best fit to the data suggesting that the connectivity patterns identified by the winning model in the value task were indeed specific to the processing of the subjective value rather than the sensory properties of the stimulus options. These findings provide compelling evidence for the existence of modality-specific communication pathways which convey value-related information across the brain.

In order to understand how reward value modulates the communication of information across the brain areas, we next examined the strength of modulatory connections in the winning model of the value task. We found that during both intra-modal and inter-modal trials, all modality-specific connections were significantly modulated at a posterior probability *P* > 0.99 (**Figure 4C**). Notably, we identified a clear distinction between the weights of feedforward and feedback connections. The directed feedforward connections from sensory ROIs to OFC ROIs and from OFC ROIs to vmPFC exhibited negative connection weights, signifying inhibitory modulatory connections. In contrast, the feedback connections displayed positive weights, indicating excitatory modulatory connections (For parameters of intrinsic connections and driving inputs, refer to **Supplementary Information: Table S4-S5**).

Additionally, to test whether effective connectivity between any two nodes of the network was better explained by unidirectional connections, we estimated all possible unidirectional variants of the winning model (i.e., model 6 shown in **Figure 4C**). As a modulatory connection between two nodes can exist in three possible ways: directed, reciprocal, bidirectional, the total number of all possible unidirectional models for the model 6 of the value task were 9, shown in **Figure 5A**. The most likely model among these was the network with bidirectional connectivity between the nodes (see the model exceedance probabilities, **Figure 5B**).

**Figure 5.**
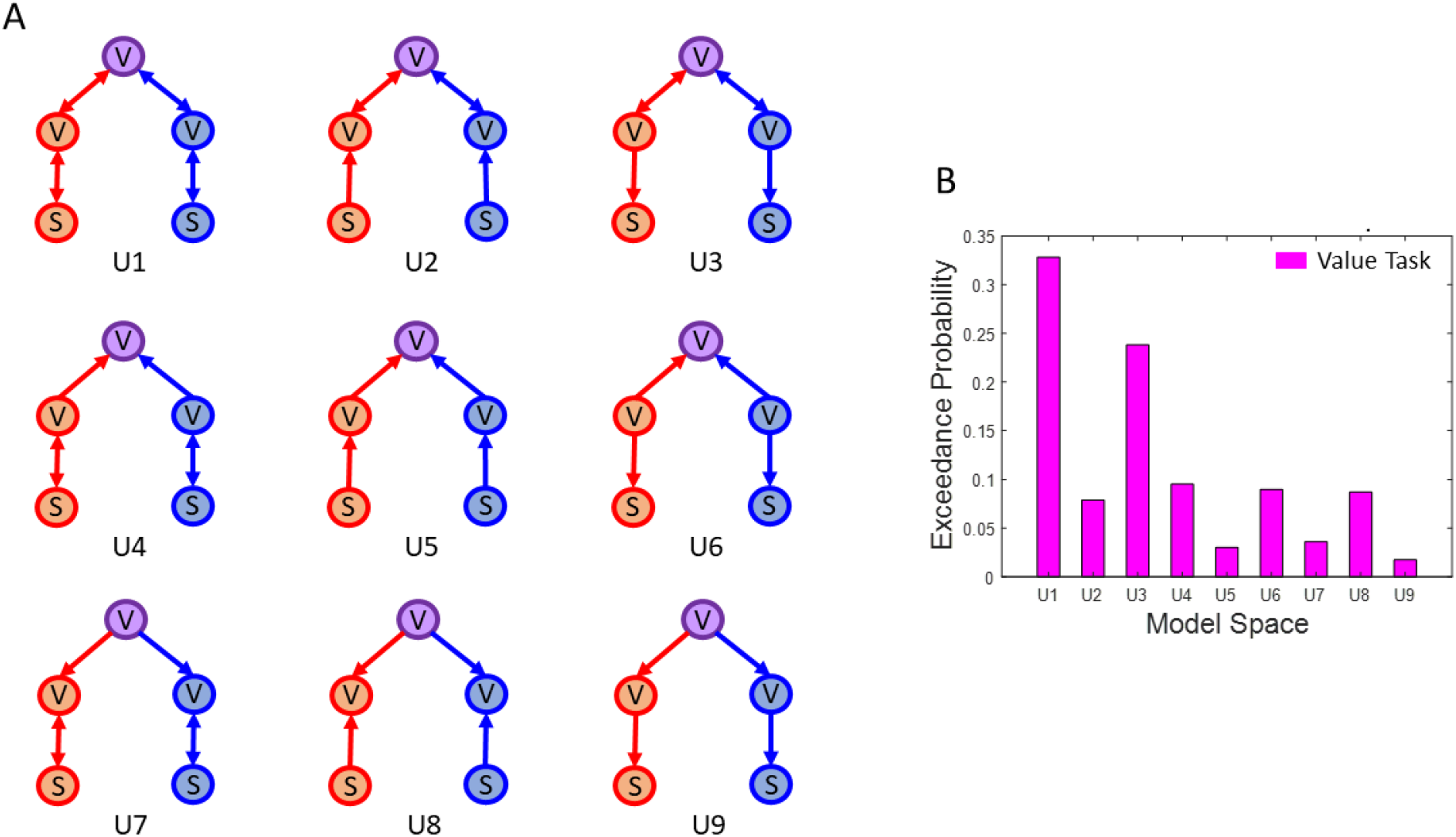
Uni- and bi-directional variants of the winning model in the value task. (**A**) Model space consisting of all possible unidirectional variants of the winning model in the value task (i.e., U1, also refer to the legend of Figure 4A and the schema in Figure 4C for information on nodes and modulatory connections). (**B**) Exceedance probabilities for the model space in (A). Model U1, i.e., a model containing bidirectional connections, is the most likely model.

Collectively, the effective connectivity results showed that auditory and visual sensory cortices communicate with separate clusters in OFC, which contain modality-specific stimulus value representations (SVR) corresponding to each sensory modality. Further, the modality-specific SVRs in OFC were linked with the modality-general SVR in vmPFC to guide the final choice.

## Discussion

In order to generate specific predictive signals for adaptive goal-directed choices, the brain must encode information about the sensory modality of reward-predicting stimuli as well as the most recent value associations with the stimuli. Moreover, to be able to compare and choose between stimuli with fundamentally distinct sensory features, general value representations are equally important. Here, we used stimuli from auditory and visual sensory modalities as reward-predicting cues in a value-based decision-making context with a dynamic foraging paradigm, enabling us to identify modality-general and modality-specific value representations using univariate fMRI analysis and reveal the underlying neural mechanisms that generate the two types of representations using effective connectivity analyses.

We found trial-by-trial value representations of auditory and visual sensory modalities to be present in segregated lateral and posterior regions of OFC, respectively. This effect cannot be due to differences in reward sensitivity or difficulty of choices since neither the choice ratios relative to rewards nor the reaction times indicated a difference between auditory and visual conditions. Examination of the generalization of this effect however, pointed towards some degree of coactivation between the two types of value representations, since tracking the auditory value in the inter-modal condition also activated the visual value representations (**Figure 3G**), perhaps due to co-activation by visually presented placeholders. Furthermore, we verified using the control task that the segregation in modality-specific representations is not due to the differences in sensory processing mechanisms underlying the auditory and visual sensory modalities. Thus, our findings show for the first time that dedicated neuronal populations exist in OFC that reflect updates in value associations of a particular sensory modality and generate specific predictive signals. As such, the present findings are in line with the known role of the OFC in representing a “cognitive-map” of the task space (Stalnaker et al., 2015; Wilson et al., 2014), especially when a task involves reversal learning (Stalnaker et al., 2015; Tsuchida et al., 2010) or devaluation of previously valuable options (Pickens et al., 2003). More specifically, these findings support recent proposals that the representation of stimulus value is an active hierarchical process in which the OFC plays a key role in representing the value of individual features of a stimuli (Hunt et al., 2014; O’Doherty et al., 2021).

In contrast to the modality-specific value representations found in OFC, we found modality-general value representations in vmPFC. Specifically, auditory value representations were found to overlap with visual value representations, aligning with the concept of the vmPFC as a common currency coding hub for distinct categories of rewarding stimuli (Chib et al., 2009; Hare et al., 2011, 2010, 2008; Lebreton et al., 2009; Levy and Glimcher, 2011; Lin et al., 2012; McNamee et al., 2013; Rolls et al., 2010; Smith et al., 2010). However, subdivisions from general to specific valuation have been also shown in vmPFC in the anterior-to-posterior direction (Clithero and Rangel, 2013; Kringelbach and Rolls, 2004; McNamee et al., 2013; Sescousse et al., 2010; Smith et al., 2010), where anterior vmPFC represents values of distinct reward categories in a general manner and posterior vmPFC in a specific manner. The loci of overlapping activations that we found in this study were in anterior vmPFC and thus support a role of the anterior vmPFC in common currency coding of value. However, we did not find any modality-specific value representations in (posterior) vmPFC, which may either be due to OFC being exclusively responsible for implementing modality-specificity in a task such as ours, or be related to the specific type of reward category, i.e., monetary rewards, that we employed in our task (McNamee et al., 2013). Interestingly, we also found that vmPFC activations were present in the control task where no continuous and gradual value-related information processing was needed. This observation points to a general role of vmPFC in computation of choice in addition to valuation, as suggested by recent theoretical frameworks (Klein-Flügge et al., 2022).

Whereas the majority of previous studies have underscored a common currency coding of subjective value, few recent studies have provided evidence for the stimulus-specific representations of reward-predicting stimuli (within visual modality) or reward outcomes (Howard et al., 2015; Howard and Kahnt, 2017; Klein-Flügge et al., 2013; Lim et al., 2013; McNamee et al., 2013), as well as modality-specific valence (Čeko et al., 2022). We are only aware of one recent human imaging study (Shuster and Levy, 2018), where modality-specificity of reward-predicting valuation across auditory and visual domains was examined. Using a risk-evaluation task where monetary values of lotteries were either presented visually or announced aurally, this study found that the anterior portion of vmPFC represents value irrespective of the sensory-modality, whereas no evidence for modality-specificity was found beyond sensory areas. However, in this study, the task design rendered the modality-specific value information redundant. As both visual and auditory stimuli could be directly translated to the same abstract numeric value and past reward history was uninformative of the current expected reward, valuation process in their task could occur entirely without tracking the sensory modality of options (Shuster and Levy, 2018). In contrast, in the present study, trial-by-trial tracking and updating of computed values of specific sensory stimuli was necessary, thus allowing us to tap into the intricate role of OFC in modality-specific updating of value. Another important aspect of our approach was to account for the covariation of sensory features and reward value (Howard and Kahnt, 2021) in determining the neural responses. This was done by employing a control task that was identical to the value task in terms of sensory requirements and final choice but differed in whether updating of computed value of each sensory modality was necessary or not. Together, a dynamic reward structure and the comparison against a task with different dimension of decision variable allowed us to unravel the co-existence of modality-specific and modality-general representations in the frontal cortex.

Apart from the frontal cortex, we found value modulations in sensory cortices, which provide evidence that representations of value are not restricted only to higher cognitive areas, as has been shown before (Serences, 2008). The value representations in sensory cortices were largely modality-specific, which means that individual sensory cortices represented the value of stimuli presented in their own sensory domain, as has been previously shown for the representation of value (Shuster and Levy, 2018) and valence (Čeko et al., 2022). These findings raised interesting questions regarding whether and how a communication of value-related information exists between sensory cortices and valuation regions. Interestingly, we found that the auditory and visual sensory cortices were bi-directionally connected to the lateral and posterior OFC (corresponding to auditory and visual value representations), respectively, in a modality-specific manner. Specifically, the modality-specific effective connectivity results revealed a high degree of selectivity: in trials when planning to choose auditory reward stimulus, there was a significant connectivity from the auditory sensory cortex to lateral OFC for these trials and not otherwise. A similar modality-specific significant connectivity existed from visual sensory cortex to posterior OFC when choosing visual reward-predicting stimulus. This finding is in line with a previous work showing connectivity between OFC and piriform cortex (relevant in case of odour stimuli) for the formation of stimulus-specific value representations in OFC (Howard et al., 2015). Moreover, past studies have shown that lateral and posterior regions of OFC receive direct afferent inputs from auditory and visual sensory cortices (Barbas, 1993; Carmichael and Price, 1995), providing neuroanatomical support for our findings.

The connectivity between sensory cortices and modality-specific representations in the OFC allows the frontal valuation areas to have access to sensory representations of rewarding stimuli in each sensory area. The sign of this connectivity provides additional insights into how modality-specific valuation is implemented. We found that feedforward connectivity in modality-specific networks was inhibitory. Feedforward inhibition has been suggested as a key mechanism in imposing temporal structure to neuronal responses (Womelsdorf and Everling, 2015) and expanding their dynamic range of activity (Pouille et al., 2009). These mechanisms allow the OFC to form an integrated representation of the sensory and value information across time, rather than encoding the precise sensory features of stimuli at every moment (akin to sensory cortices), thereby serving as a cognitive temporal map of the task (Stalnaker et al., 2015; Wilson et al., 2014). Additionally, connectivity results showed that the value modulations in sensory cortices were driven by top-down feedback signals generated in respective valuation regions in OFC. This is consistent with previous work showing that biasing signals generated from frontal and parietal areas modulate spatially selective visual areas (Serences, 2008). In fact, recent studies have provided robust causal evidence for the role of lateral OFC in value-driven guidance of information processing in sensory cortices (Banerjee et al., 2020). Our finding of the presence of excitatory feedback connectivity between the modality-specific representations in lateral and posterior OFC and auditory and visual cortices, provides strong support for the causal role of top-down valuation signals in shaping sensory perception during decision-making.

Further, we found that specific value representation in OFC were linked to general value representations in vmPFC. Specifically, we showed that when planning to select an auditory reward-predicting option, there was a change in the connectivity between the auditory value representations in OFC and modality-general representations in vmPFC, with a similar pattern found for the selection of the visual options. This result highlights the underlying mechanism whereby value representations in OFC provide input to the vmPFC to support the formation of general value representations needed for the comparison of options from distinct modalities and deriving the final choice. Notably, the modality-specific connectivity between OFC and vmPFC corroborates previous findings showing that sensory-specific satiety-related changes in connectivity between OFC and vmPFC predicted choices in a devaluation task (Howard and Kahnt, 2017). Together, these results show how common currency coding of value integrates modality-specific information about reward-predicting stimuli in a dynamic environment to guide choices.

Understanding whether valuation signals in frontal cortex contain information about the sensory modality of reward-predicting stimuli has a number of important theoretical and clinical implications that go beyond the specialized field of neuroeconomics and value-based decision making (Levy and Glimcher, 2012; Padoa-Schioppa and Schoenbaum, 2015; Rangel et al., 2008). We show that value-based choices involving reward-predicting stimuli from different sensory modalities are supported by bi-directional connectivity between the sensory areas and the modality-specific representations in OFC. Although top-down modulation of perception through interactions between frontal and sensory areas has been the basic tenet of a number of influential theoretical frameworks (Corbetta and Shulman, 2002; Desimone and Duncan, 1995; Friston, 2005; Gardner and Schoenbaum, 2021), the importance of modality-specific representations of reward value in frontal areas that could provide a biologically plausible implementation of these putative interactions has been largely ignored. Therefore, our study provides novel insight for future computational work on how top-down signals can be selectively routed to impact on sensory processing. In doing so, it is important to note that the modality-specific representations that we found may adapt and reorganize under different contexts rather than being hardwired and fixed in the brain (see also Antono et al., 2023). In fact, outcome-related adaptation in the representation of value can occur during the same task (Rich and Wallis, 2016), which provides a flexible mechanism for reorganizing neuronal codes of value based on the context. Future studies will be needed to examine whether and to what extent the modality-specific coding of value can adapt to the specific features of a task. From a clinical perspective, our results suggest that localized lesions to OFC may be associated with specialized impairments of value-based decisions in visual or auditory domains, an interesting possibility that can be further investigated by future studies. Additionally, our findings may allow a better understanding of pathological states such hallucinations (Frith, 1996; Rolls et al., 2008) where illusory percepts arise in the absence of external stimuli (Powers et al., 2016), likely due to the aberrations in communication pathways between the frontal and sensory areas (Allen et al., 2008). More generally, the present study, together with previous efforts in understanding how value-related information is communicated between the frontal and sensory areas (Banerjee et al., 2020; Howard et al., 2015; Howard and Kahnt, 2018), provide instrumental insights regarding how perceptual and cognitive processes are coordinated in the brain.

In summary, our results provide novel evidence for the co-existence of modality-specific and modality-general encoding of subjective value in OFC and vmPFC, respectively, pointing to the specialized functions of these two valuation areas. A general value signal would facilitate the comparison between distinct rewarding stimuli (Levy and Glimcher, 2012, 2011) and the transformation of stimulus values into motor commands (Hare et al., 2011). On the contrary, modality-specific value encoding associated to respective sensory cortical representations would support goal-directed adaptive behaviour by generating specific predictive signals about impending goals (Nogueira et al., 2017; Stalnaker et al., 2014; Wilson et al., 2014), such as when planning to choose auditory or visual reward-predicting stimuli. We further show how the communication between sensory areas and modality-specific representations of subjective value in OFC plays a central role in supporting value-based decisions in a multimodal dynamic environment.

## Material and Methods

### Participants

Twenty-four healthy subjects (13 male and 11 female, age 19 to 45 years; mean ± SD age = 27.92 ± 6.04 years, including one of the co-authors) participated in the experiment for financial compensation of 8€/hour. The sample size was based on a previous study that used a similar paradigm (Serences, 2008). The experiment was done in two sessions, each lasting about 150 minutes (comprising 45 min preparation time and 105 min scanning time: 90 min functional and 15 min structural scans). Before the first session an online training (30 min) was scheduled to familiarise participants with the task. Participants also had the opportunity to earn a monetary bonus of maximum 22€ based on their behavioural performance in the value-based decision-making task (value task) during the scanning session. All participants were right-handed and had normal or corrected-to-normal vision. Before the experiment started and after all procedures were explained, participants gave an informed written consent and participated in a practice session. The study was approved by the local ethics committee of the “Universitätsmedizin Göttingen” (UMG), under the proposal number 15/7/15.

Four participants were excluded from the final analysis resulting in the data from 20 subjects presented here (10 male and 10 female, age 21 to 42 years; mean ± SD age = 29.00 ± 6.34 years): two participants had difficulty in differentiating the strategies of the value and the control task (specifically with the instructions associated with the feedback colours in the two tasks, see the Experimental Design); one participant was excluded due to excessive head motion while scanning (> 4 mm); and one participant due to the unusually large size of the ventricles in the structural MRI scan.

### Experimental design

The experiment consisted of a value-based (value task) and an instruction-based (control task) decision-making task, completed in two sessions (**Figure 1**). Each session consisted of 12 blocks (of 72 trials each): 9 blocks of the value task (i.e., 3 blocks for each of the 3 reward ratios; 1:3, 1:1, 3:1, for details see the *Dynamic Reward Structure*) followed by 3 blocks of the control task. Each task involved a binary choice between stimuli presented in three sensory domains: both auditory (*AudAud*), both visual (*VisVis*), and audio-visual (*AudVis*), which were presented in separate blocks. All three types of sensory domain blocks appeared an equal number of times across each task in a pseudo-random order.

#### Stimuli

Two pure auditory tones (low pitch (LP) tone-sawtooth, 294 Hz; high pitch (HP) tone-sinusoidal, 1000 Hz, played through MR-compatible earphones -Sensimetric S15, Sensimetrics Corporation, Gloucester, MA-with an eartip -Comply™ Foam Canal Tips-) and two contrast reversing visual checkerboards (green and black or red and black with the contrast reversing at 8 Hz, as in (Serences, 2008)) within circular apertures (4 ° radius) were used as the choice options. In an auditory (*AudAud*) or a visual (*VisVis*) trial, either two tones or two checkerboards were presented as options, respectively. In an audio-visual (*AudVis*) trial, one tone and one checkerboard were presented as options. Choice options were presented simultaneously on the left or right side of the centre (auditory stimuli were played one on each side of the earphones; visual stimuli were centred at the 10 ° horizontal distance and 5 ° above the centre of the screen). Different tones and coloured checkerboards and their combinations (in *AudVis* blocks) were presented an equal number of times across the 72 trials a of a block and in a pseudorandom order.

#### Trial Structure

Both the value and control tasks featured identical presentations of stimulus options, response requirements, and feedback on the decisions. The only distinction between them lay in the cues associated with the feedback colours (**Figure 1A-B**). Participants were asked to fixate continuously throughout each run (here, a run = 3 blocks) on a small square (0.4 ° visual degree) at the centre of the screen. A trial began with a mean fixation period of 1.8 s (±0.45 s), yielding a mean trial duration of 4.3 s. Following the fixation period, the two stimuli options were presented simultaneously for 1 s, one on each side of fixation. The spatial position of each option was also pseudo-randomised across the trials of a block in such a way that each option appeared an equal number of times on both sides of the fixation point. Following the onset of the stimulus options, participants pressed either the left or the right button on a MR-compatible two-buttoned response box (Current Designs Inc., Philadelphia, PA), using the index or the middle finger of their right hand, to indicate their choice. The participants were required to respond within 2.25 s following the onset of the options. Following the response window, a feedback window of 0.25 s appeared in which the central fixation point turned either yellow or blue in colour. In the value task, the yellow fixation indicated that the choice was rewarded, and the blue fixation indicated that the choice was not rewarded. Since the control task was designed to be similar to the value task in terms of sensory processing requirements without a need to track and update their estimation of options’ value, the feedback instructed the participant to make a prescribed choice. Thus, in the control task, the yellow fixation indicated to switch from the past trial choice and the blue fixation indicated to keep the past trial choice. The choice on the first trial of the control task in each block was a random choice.

Note that in all intervals during a trial in both tasks, two placeholders (circular apertures: 4 ° radius) containing white noise were presented on the screen. This was specifically done to reduce the visual after-effects which could ensue from the presentation of the coloured checkerboards, and additionally minimized the low-level sensory differences between the different stimulus configurations. When choice options were from the visual modality, the placeholders were replaced with the coloured checkerboards and otherwise stayed in view on the screen (**Figure 1A**).

#### Dynamic reward structure

To create a dynamic multimodal environment for participants, rewards were assigned to the options from different sensory modalities independently and stochastically at random intervals using a Poisson process (Corrado et al., 2005). On average, a reward was available for delivery on 33% of the trials (of a block of value task). These 24 rewards in a block (33% of 72 trials) were distributed between the two stimuli options in different reward ratios of {1:3, 1:1, 3:1}, such that the rewards assigned to options were {8.5%:24.5%, 16.5%:16.5%, 24.5%:8.5%} in percentage of trials. For the value task in a single session (9 blocks), these three reward ratios were repeated and randomised such that each reward ratio was used exactly once with every sensory domain block (i.e., *AudAud*, *VisVis*, and *AudVis*). The randomisation of various factors such as sensory modality, spatial position of options, and reward ratios was done to provide a dynamic environment, in which the participant would be required not only to update their stimuli-value associations with changing reward ratios but also to keep track of the reward-predicting stimuli very carefully on a trial-to-trial basis as spatial positions changed. Two important schemes of “baiting” and “change over delay” (COD) were adopted as in previous studies (Corrado et al., 2005; Serences, 2008). In baiting, an assigned reward to an option remained available until that option was chosen. This was done to avoid the “extreme exploitation” strategy in which a participant would always stick to the option with a higher reward rate (e.g., 24.5%>8.5%) association in a block and to motivate the exploration in which a participant should visit both options occasionally. Also, an earned reward feedback was delayed for one trial when the participant changed their choice from one option to the other and delivered only if the participant chose the same option again. This cost, i.e., COD, was employed to discourage “extreme exploration” strategy, where the participant would be able to consume all rewards without any learning by alternating choices rapidly between options. Trials following a change of choice (switch) between options were not included in the behavioural analysis (and were marked by a specific regressor in fMRI analysis) because subjects were informed that they will not get a reward on such trials and hence choices were not completely free. At the end of each block, participants were shown the reward earned in that block at the rate of 5 cents per yellow square shown as the reward feedback. At the end of the second session, participants received the total reward earned which was up to a maximum of 22€ (11€ per session) based on their performance along with a participation fee of 8€ per hour.

#### Control task structure

Similar to the reward structure in the value task, switches were assigned independently and stochastically to the options in an equiprobable manner with an average switch rate of 33%. Thus, on any trial when a participant earned a switch from a chosen option, yellow feedback was displayed indicating that they should switch their choice to the other option on the next trial. On other trials, when a switch was not assigned, blue feedback was shown to indicate that the same option should be chosen on the next trial. This type of switch assignment structure was developed to encourage a similar temporal choice pattern as in the value task. On a single day, the control task was conducted in each of the three sensory domains. There were no baiting and COD schemes employed in the control task. At the end of each block, participants were shown their performance that indicated how accurately they followed the instructions in that block.

### Computational framework of choice behaviour

To examine whether participants’ choices in the value task were influenced by the dynamic reward structure employed in our design, we used a computational framework that has been used in the past to model choice behaviour abiding by the matching law (Corrado et al., 2005; Herrnstein and Baum, 1970; Serences, 2008). In our task, there were no prior reward associations with the options, and hence on any trial *t* a participant made a choice *c*(*t*) based on the previous rewards received *r*(*t*^-^) during the experiment, see **Figure 1C**. Intuitively, an option that delivers more rewards per unit of time should have relatively higher value associations and should be chosen more often. Thus, to estimate participants’ subjective value beliefs for each reward option on a trial-by-trial basis, we fitted the reward history and choice data of each participant to a linear-nonlinear-probabilistic (LNP) model, shown in **Figure 1C** (also called as linear regression-based model of reinforcement learning (Katahira, 2015)). We chose an LNP model over other reinforcement learning models (RL models such as Q-learning) since the former has been shown to better capture the statistics of the matching behaviour that is observed in our paradigm (Corrado et al., 2005). Two broad phases of the LNP model are the *learning* and the *decision-making* phase (for implementation details refer to Corrado et al., 2005; Serences, 2008).

In the *learning phase* (see **Figure 1C**), two identical linear filters (*n* learning weights *α*_*τ*_, *τ* = 1 to *n* trials in the past) weigh the reward history (till *n* past trials) of each option (*r*_i_(*t*^-^), *i* = 1,2 corresponding to stimulus sets *S*_1_, *S*_2_), where *n* is equal to half of the trials over which the reward ratio was unchanged (here, *n* = 36) (Serences, 2008). The purpose of the filter is to look back on the reward history of past *n* trials and distil the effect of those trials into a scalar value. This scalar value should be representative of the participant’s expectations for associated value of stimulus options. In other words, if the choice of a particular option was rewarded (or not rewarded) on the past trials, then the value belief for that option should be relatively higher (or lower) on the current trial, respectively.

First, we explain how filter weights were derived from the reward history and choice data of a participant and then how these weights were used to compute the subjective value beliefs for the stimuli options on each trial. The reward history *r*_i_(*t*^-^) of a stimulus option *S*_i_ is a binary vector containing 1 when that option was chosen and rewarded on trial *t* and 0 when chosen and not rewarded on trial *t* or simply when not chosen on that trial. As explained in previous section on experimental design, trials following a change of choice (switch) between options were not included in the behavioural analysis because subjects were informed that they will not get a reward on such trials and hence choices were not completely free. Thus, the free choice *c*_*i*_(*t*) of a stimulus option *S*_*i*_ is denoted by 1 when that option was chosen on trial *t* and 0 when not chosen on that trial. As the overall reward assignment over the two options was symmetric, their impact on choice was equal and opposite, hence we used the composite reward history ***r*** (as shown in (1)) and composite choices ***c*** (shown in (2)) for further analysis.

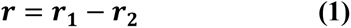

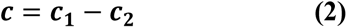

A participant’s choices are strongly influenced by the reward history (the learning mechanism), thus, intuitively the filter weights should most closely match the input reward history to the desired output choices. This is an optimization problem (Corrado et al., 2005; Serences, 2008) and can be solved by employing Wiener-Hopf equations:

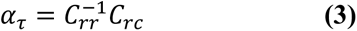

where, *C_rr_* is auto-covariance matrix of input time series *r*(*t*^-^) and *C_rc_* is cross-covariance vector of output time series *c*(*t*). Further, what we used in the analysis were relative filter weights α̂_τ_, which were obtained by normalising weights obtained in (3) such that the sum of all *n* weights equals 1 (**Figure 2C** shows relative filter weights of a single participant). For more implementation details refer to (Corrado et al., 2005; Serences, 2008).

Next, the relative filter weights were used to compute subjective value beliefs for the stimuli options *v_i_*(*t*), *i* = 1,2 on trial *t*. This was done by multiplying the reward history of option *i* on past *n* trials by the weighting coefficients α̂_τ_ and summing the product results over the past *n* trials (see **Figure 1C**):

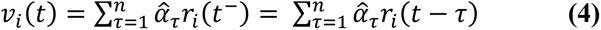

The *decision-making phase* (see **Figure 1C**), draws the ultimate binary choice (*S*_1_ or *S*_2_) on trial *t* based on a relation that maps the differential value *dv*(*t*) (as shown in (5)) computed on trial *t* to the participant’s probability of choosing option *S*_1_on that trial. Intuitively, this relation should strongly predict a participant’s choice behaviour, where the participant should make a choice *c*(*t*) based on the comparison process shown in (6).

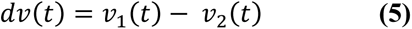

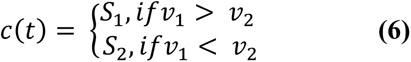

To assess the fit of the LNP model during the *learning phase*, the relative filter weights for the data of each participant were approximated by fitting an exponentially decaying function, indicating that choices were most impacted by recent rewards rather than distant rewards in past (quantified by time scale parameter *τ* trials of the fit; see **Figure 1C** for the illustration of the filter and **Figure 2C** for the fit to the data of a single participant). To assess the *decision-making phase*, the probability of choosing option *S*_1_ for each participant was approximated by fitting a normal cumulative distribution function (equation (7)).

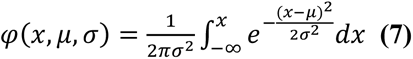

where x is the differential value (*dv*). This function contains two important decision-making parameters: *µ* corresponding to a participant’s biasness towards a particular option and *σ* that measures the sensitivity to value differences or in other words the explore-exploit tendency. Accordingly, *σ* = 0 corresponds to an extreme exploitative tendency, and *σ* → ∞ to extreme exploration. The disadvantage of being extremely exploitative; i.e., sticking to an option that has higher reward rate associated with it, is that it would yield lesser number of rewards to the participant because there exist unvisited options, which remain baited until chosen. Moreover, extreme exploration would also be disadvantageous, as it would lead to no learning and the absence of any strategy. Thus, the optimal strategy in this task would be to choose more often the option with higher reward rate and to occasionally visit the less rewarding option to consume rewards on it. An optimal strategy is advantageous in a dynamic reward structure task where the aim is to maximize rewards, and to examine whether this is the case in our task, we inspected the abovementioned parameters (*τ*, *µ*, *σ*) for their fit to participants’ behavioural data (**Figure 2C-E**).

In the value task, the positive and negative feedbacks have distinct effects on participants’ beliefs. Therefore, if the choice of a particular option was rewarded (yellow feedback) or not rewarded (blue feedback) on the previous trial, then the value beliefs for that option should be relatively higher or lower, respectively, on the current trial in comparison to the value beliefs in the past trial. As only one of the two options could be chosen and rewarded in any trial, the differential value of two options would also be relatively high in magnitude when reward was received on the past trial, and otherwise low. On the contrary, the control task was designed in a way to be like the value task in terms of sensory processing requirements but not involve the participant in any learning or updating of the stimuli value. Intuitively, when no learning via feedbacks occurs, the two types of feedbacks (keep/switch) should have a similar effect on the subjective preference over the options. To confirm this, we tested the fit of the same LNP model to the choices in the control task and compared the absolute differential values of each trial obtained from models’ fits to both tasks (value and control tasks) against the type of feedback received (blue or yellow) in the previous trial (**Figure 2F**). We used the absolute differential values (*absDVs*) as a measure of subjective preferences because the choice behaviour is symmetric with respect to the individual options.

### fMRI data acquisition and pre-processing

MRI scanning was carried out on a 3-Tesla Siemens MAGNETOM Prisma scanner equipped with a 64-channel head-neck coil at the Universitätsmedizin Göttingen. Anatomical images were acquired using an MPRAGE T1-weighted sequence that yielded images with a 1 x 1 x 1 mm resolution. Whole-brain multi-shot Echoplanar imaging (EPI) volumes were acquired in 69 interleaved transverse slices (TR = 1500 ms, TE = 30 ms, flip angle = 70°, image matrix = 104 x 104, field of view = 210 mm, slice thickness = 2 mm, 0.2 mm gap, PE acceleration factor = 3, GRAPPA factor = 2). Data from each participant was collected in two identical sessions on two separate days. An experimental session consisted of multiple runs of fMRI data acquisition, where a run comprised starting the scan and acquiring data for three blocks of the tasks (∼ 20 minutes) after which the scan was stopped and resumed again after a break (∼ five minutes). On each day, four fMRI runs (first three runs: 9 blocks of the value task, last run: 3 blocks of the control task) were conducted and each fMRI run lasted 16.355 min.

Data pre-processing and further statistical analyses were performed using Statistical Parametric Mapping software (version SPM12: v7487; https://www.fil.ion.ucl.ac.uk/spm/) and custom time-series analysis routines written in MATLAB. EPI images of each session were slice time corrected, motion corrected, and distortion corrected by using the measured field maps. The T1 anatomical image was co-registered to the mean EPI from realign-&-unwarp step, and then segmented. The estimated deformation fields from the segmentation were used for spatial normalisation of the corrected functional and anatomical images from the native to the MNI (Montreal Neurological Institute) space. Finally, the normalised EPI images were spatially smoothed using a 6 x 6 x 6 mm FWHM Gaussian kernel.

### fMRI univariate analysis: General linear modelling (univarGLM)

For each participant, we first specified a general linear model (GLM) using the pre-processed functional images of two sessions that were concatenated. The GLM modelled both the value and the control task using 35 event-related regressors convolved with the canonical hemodynamic response function (HRF). For the value task, we defined individually for each of the three modality conditions (auditory, visual, audio-visual) one unmodulated stick regressor representing the modality-wise trial identity and two parametrically modulated stick regressors containing the trial-by-trial updated subjective value (SV) beliefs regarding each of the options presented, referred to as the value-modulated regressors. Trial identity was entered as 1 at the onset of the stimuli for trials of a particular modality condition and 0 otherwise. The value-modulated parametric regressors (SVs) were estimated based on the LNP model (see **equation 4**) and represented the trial-by-trial learning and updating of subjective value of each option separately (**Figure 1C** and **Supplementary Figure S3**). The trial-by-trial SVs were entered at the onset of the stimuli options and are denoted by *lpSV* and *hpSV* in auditory domain corresponding to low pitch and high pitch tones, *rSV* and *gSV* in visual domain corresponding to red and green checkerboard stimuli, *aSV* and *vSV* in audio-visual domain corresponding to auditory and visual stimuli in any combination (see also **Table 1**).

Similarly, for the control task we defined individually for each of the three modality-domains one unmodulated regressor representing the modality-wise trial identity and two parametrically modulated regressors corresponding to each of the options presented. In the control task, the aim was to passively follow instructions. Thus, to create a parametrically modulated regressor corresponding to one stimulus option, a weight of either 1 or 0 was assigned at the onset of stimuli options in each trial depending on whether the instruction (keep/switch your choice) from the last trial was correctly followed or not, respectively (see also **Supplementary Table S1** and **Figure S3**).

In order to separate signals related to the expected value of the stimuli from the signals related to the choice (response) and receipt of an outcome (feedback), we included two unmodulated event-related regressors (collapsed across the value and the control task) locked to the time of response and the onset of feedback in our GLM. Note that in our paradigm, in any current trial ‘*t*’, the stimuli-value associations are updated based on the feedback event in the previous trial ‘*t*-1’. Therefore, in our task design we had introduced a jitter between the feedback in the previous trial and the stimuli presentation in the current trial. This jitter together with modelling the feedback and response with separate repressors allowed us to separate the signals related to the expectation of stimuli value from the signals related to the receipt of an outcome, which would have been otherwise impossible due to the sluggish nature of the hemodynamic responses. Furthermore, 15 nuisance regressors were included in the GLM corresponding to the following: instruction presentation at the start of each block, six motion parameters, run regressors (modelled by assigning a weight of 1 for each volume of that run and else 0: a run corresponds to each period of MRI data acquisition between the start and the end of the scan) to account for the difference in the mean signal activity between each time the scan started (one less in number than the total number of fMRI runs, here 7) and a constant.

Based on the GLM described above, the brain regions which represented subjective value in each domain were identified by inspecting several univariate contrasts of the value-modulated parametric regressors (**Figure 3** and **Table 1-3**). Note that our primary interest in this study was to identify the neural correlates of valuation for different sensory configurations. Since in a binary choice situation valuation occurs for each of the two options separately, we used the trial-by-trial value-modulations of each option as an independent variable (one parametric regressor for each option in our GLM) to identify areas in the brain which encode the value of each option. This is a different approach than using the differential value of the options as the independent variable which seeks to identify the neural correlates of comparison and choice between options (Serences, 2008) rather than the valuation of each individual option which occurs before the choice process. Accordingly, in intra-modal conditions (*AudAud* and *VisVis*), we estimated an overall effect of value-modulated regressors separately in auditory and visual sensory domains by defining contrasts: *intraAudSV* > 0 and *intraVisSV* > 0. The auditory contrast *intraAudSV>0* is the overall effect of parametric regressors of interest *lpSV and hpSV* (according to SPM convention, the contrast vector corresponding to 35 regressors contains 1 for the two parametric regressors of interest, here *lpSV* and *hpSV*, and 0 otherwise). Similarly, for the visual contrast *intraVisSV>0* is the overall effect of parametric regressors *rSV and gSV.* In the inter-modal condition (*AudVis*), we estimated the overall effect of value modulations in both sensory modalities using the contrast: *interAudVisSV* > 0, where *interAudVisSV>0* is the overall effect of parametric regressors of interest *aSV and vSV*.

On account of previous studies identifying the domain-general and domain-specific valuation areas in the vmPFC and OFC (Howard et al., 2015; Howard and Kahnt, 2017; McNamee et al., 2013), we limited our analysis to a mask encompassing the orbital surface of frontal gyrus. This search volume (for details see **Figure S2**) consisted of anatomical parcellations of orbital surface of frontal gyrus as defined in automated anatomical labelling (AAL) atlas (Rolls et al., 2020, 2015). Statistical maps were assessed for cluster-wise significance using a cluster-defining threshold of *t*(19) = 3.58, *P* = 0.001 for simple contrasts (see **Table 1**) and at *t*(19) = 2.86, P = 0.005 for interaction contrasts (see **Table 2**); and using small volume corrected threshold of P < 0.005 (referred to as a small volume family-wise-error (SVFWE) correction) within the frontal search volume. Whole-brain results were inspected at FWE p<0.05, and k>10 (see **Table 3**).

To perform a cross-validation of our results regarding the modality-specificity of the subjective value representations in the lateral OFC, we defined ROIs based on the data of intra-modal condition and tested the generalization of our results to the data of inter-modal condition (**Figure 3G**). For this, the estimated effect size of the parametric value regressors (t-value) in each ROI (red and blue clusters in **Figure 3A-E**) was used to define a modality-specificity index (i.e., the difference between responses to the same versus different modalities with respect to each ROI). Based on this index, the 50 most modality-specific voxels of each ROI were selected (∼20% of the voxels contained in red and blue clusters of **Figure 3A-E**). We then tested the extent to which the modality-specificity in these voxels generalized to the data of inter-modal condition.

### Effective connectivity analysis of fMRI data

Our univariate analysis identified a number of regions, both in sensory and in frontal areas, that were modulated by the subjective value of each choice option (**Figure 3**, **Table 3** and **Supplementary Figure S5**). We next aimed to determine how the long-range communication between these areas generates the stimulus value representations (SVRs) and the degree to which these representations are modality-specific. To this end, we investigated the effective connectivity (EC) of a network consisting of sensory and frontal regions exhibiting value modulations at the time of options’ presentation by employing deterministic bilinear dynamic causal modelling (DCM) approach (Friston, 2011; Friston et al., 2003; Stephan et al., 2009). This approach fits a set of pre-defined patterns of EC within a model space to the fMRI time series and compares them in terms of their evidence (for details of the model space see the section under *Defining the model space for a 5-node network*).

The DCM approach requires two basic types of information for extracting time series data: the regions of interest (ROI, e.g., *audOFC,* see **Figure 4** and the section under the **Regions of interest**) and the onset times of experimental conditions (e.g., intra-modal auditory condition of value task). The predefined model space contains the ROI information and the information on different experimental conditions can be provided by a GLM containing trial onsets of the experimental conditions that are of interest. The GLM used for effective connectivity analysis contained four unmodulated choice identity regressors. Two of these regressors represented the final choice (whether auditory or visual stimulus) for each intra-modal condition (*AudAud* or *VisVis*). The other two regressors corresponded to the inter-modal condition, separating these into auditory and visual trials based on the final choice (whether auditory or visual stimulus). Note that in intra-modal conditions (*AudAud*, *VisVis*) effective connectivity is the connectivity among sensory and frontal regions during the valuation of a particular modality condition, either auditory or visual. However, for inter-modal condition (*AudVis*), changes in EC during the valuation process occur for both auditory and visual modalities. To test the same models for their fit to both intra- and inter-modal conditions, we separated trials in the inter-modal condition according to whether the auditory or visual stimulus was selected (hence denoted by *interAud* or *interVis*, respectively). Using the same model space for both intra- and inter-modal conditions would provide insights into the underlying mechanisms that mediate valuation across different contexts, i.e., when the same or different sensory modalities are compared against each other in terms of their value. Additionally, this approach is more parsimonious than either having two separate sets of models for each condition or increasing the number or complexity of the models to account for the differences between intra- and inter-modal conditions (Vandekerckhove et al., 2015).

**Regions of interest (ROIs):** ROIs for the effective connectivity analysis comprised the frontal valuation areas and the sensory regions that contained stimulus value representations for auditory and visual modalities according to the univariate analysis (**Figure 3A-E**, **Table 3**, and **Supplementary Figure S5**). The resulting five ROIs from which time series for DCM analysis were extracted were as follows: 1) The overlapping activation area for intra-modal visual and auditory value representations in vmPFC, 2) The activation area of left latOFC during intra-modal auditory condition – i.e., audOFC, 3) The activation area of left postOFC during intra-modal visual condition – i.e., visOFC, 4) bilateral activations in auditory sensory cortex – i.e., audSen, and 5) bilateral activations in visual sensory cortex – i.e, visSen. From each ROI the first principal component of the pre-processed fMRI time series was extracted and fed to the EC analysis (Zeidman et al., 2019).

**Defining the model space for a 5-node network:** In order to understand how valuation is supported by a network comprising modality-general and modality-specific representations, we estimated 22 biologically plausible models for the value and the control tasks (11 models in each task). These models were developed over a base model and contained three types of connections: driving input, intrinsic, and modulatory connections. In all models, intrinsic connections were defined for every node of the network as self-connections and independent of the experimental condition (see also the **Supplementary Tables S4 and S5**). Because the stimuli were presented aurally or visually, two types of driving inputs to the network were defined for auditory and visual sensory cortices: 1) an input to ROI audSen in auditory and audio-visual conditions of both tasks, and 2) an input to ROI visSen in visual and audio-visual conditions of both tasks. The driving input was modelled by entering ones at the onset of stimuli options belonging to a certain condition type and zeros otherwise. The different models differed from each other with respect to the modulatory connections between nodes, which depended on the experimental conditions. The model space of all possible connectivity models would be extensive for a 5-node network (Friston et al., 2011), where a modulatory connection between any two nodes can exist in none or more of the 4 experimental conditions of a task (*intraAud*, *intraVis*, *interAud*, *interVis*) and 2 directions (directed and reciprocal). Therefore, we constrained the model space based on the following assumptions, which resulted in model 1 shown in **Figure 4A**:

1. We included models with only bidirectional modulatory connections between nodes (Friston et al., 2011), based on the past findings that anatomical connectivity between two cortical areas is generally bidirectional (Zeki and Shipp, 1988). Additionally, large connectivity databases indicate a strong likelihood of cortico-cortical connections to be bidirectional (Kötter and Stephan, 2003). Moreover, this constraint does not imply that connection strengths would be identical for both the directed and reciprocal connections between two nodes.
2. Based on our specific hypotheses and for simplicity, we only included models that assumed two distinct and symmetric sub-networks for the valuation of stimulus options from auditory or visual modalities (auditory modality in: intraAud and interAud; visual modality in intraVis and interVis). The auditory sub-network comprised auditory valuation (vmPFC, audOFC) and sensory (audSen) ROIs and visual subnetwork comprised the visual valuation (vmPFC, visOFC) and sensory (visSen) ROIs. Models which contained modulatory connectivity between sensory areas or between OFC clusters were not included in our model space, however intrinsic connections exist between every pair of ROIs.
3. Finally, to explicitly test the plausibility of modality-specific valuation in OFC, we focused on EC patterns in which modality-specificity was either present or absent. The presence of modality-specific EC assumes that a sensory area (e.g., audSen) should only communicate to its respective value representation in OFC (i.e., audOFC) and not to others (i.e., visOFC), during experimental conditions involving that sensory modality (i.e., in intraAud, interAud). The absence of modality-specific EC assumes that additionally there are cross-modality connections (see model 1 in **Figure 4**) between each sensory area and OFC representations. In this case, each sensory area (e.g., audSen) communicates not only to its respective OFC (i.e., audOFC) during the auditory conditions but also communicates to the other OFC value representations (i.e., visOFC) during all experimental conditions (i.e., intraAud, interAud, intraVis, interVis).

Based on the above assumptions, we first built a biologically plausible full model (model 1 in **Figure 4A**). We then constructed the model space, exhaustively, by generating models with a subset of connections from the full model. In doing so, we took care of the following aspects: 1) we maintained the subset connections in a model symmetrical across the auditory and visual subnetworks; 2) we considered cross-modality connections either for all or for none of the conditions; and 3) we included models with each node connected to at least one other node of the network. Additionally, we also included a null model, which had no modulatory connectivity in the network for any experimental condition. This approach resulted in a connectivity model space consisting of 10 models per task (shown in **Figure 4A**) plus a null model. In this model set, models 2, 6, 7, 8 assumed modality-specific EC and models 1, 3, 4, 5, 9, 10 assumed the lack of modality-specificity (i.e., the existence of cross-modality). We estimated each of the 22 models (11 in each task) individually for all 20 subjects. However, for one subject the parameter estimation did not converge and therefore, we excluded that subject from the effective connectivity analysis. Thereafter, we identified the most likely model using a group-level random effects Bayesian model selection (rfxBMS) approach (Stephan et al., 2009). The model exceedance probability used to find the best model as shown in **Figure 4B** represents the probability that a particular model *m* is more likely than any other model in the model space (comprising of *M* models), given the group data. Note that the exceedance probabilities over the model space add to one (Stephan et al., 2009). Next, we estimated the connection strength parameters for connections of interest using a Bayesian parameter averaging (BPA) approach (**Figure 4C**). After identifying the model that best characterized EC in the value task, we further confirmed that the bidirectional variant of the winning model outperformed the unidirectional variants (**Figure 5**).

## Acknowledgements

We thank Tabea Hildebrand and Jana Znaniewitz for their help with the data collection. This work was supported by an ERC Starting Grant (no: 716846) to AP.

## Authors’ contributions

SD and AP conceptualized the project and designed the main task. SD, IK and AP designed the control task. SD, JEA, and AP conducted the experiments. JEA preprocessed the data. SD and AP analysed the data. SD and AP interpreted the results and wrote the first draft of the manuscript. All authors revised the manuscript. AP acquired funding.

## Data and code availability

The datasets generated and analysed in the current study are available from the corresponding author upon request.

## Supplementary Text and Tables

### Definition of parametric regressors for the control task

Similar to the value task, two regressors were modelled at the onset of stimuli options in the control task: one unmodulated regressor representing the modality-wise trial identity and two parametrically modulated regressors for each of the two choice options (*S*_1_ and *S*_2_). To define the parametric regressors, we assigned a weight of either 1 or 0 to each option according to the schema shown in **Table S1**.

### Relationship between reward ratios in each modality and the probability of choice

We found a weak main effect of modality (F[2,38] = 5.95, p = 0.024) on choice ratios, indicating that choice ratios differed between modalities. This effect corresponded to a tendency of participants to choose the visual option more often than the auditory option in the audio-visual block even when they had the same reward ratio 1:1 as can be seen in **Figure 2A**, thereby creating a difference between choice ratios of intra- and inter-modal conditions. However, this difference only reached significance for the reward ratio 3:1 as in inter-modal trials as participants chose the auditory modality significantly less often than options in intra-modal trials (**Table S2**). Note that since LNP models fitted to the fMRI data were estimated for each condition separately, this bias (i.e., preference of visual over auditory stimuli in audiovisual blocks) did not have any impact on our reported results regarding the differences of value representations between modalities.

### Analysis of reaction times (RTs)

We conducted two analyses of RTs. Firstly, we carried out an analysis similar to the one done for choice ratios and shown in **Figure 2A-B**. The aim of the RT analysis was to test whether there was a systematic difference between the auditory and visual modalities in terms of their processing requirements, for instance the task difficulty. As shown in **Figure S1**, we found that intermodal responses were systematically slower than intra-modal responses F[2,38] = 12.4, p <0.001, but importantly there was neither a significant effect of reward ratio nor interaction with modality (Fs<1 and ps>0.1), and RTs of auditory and visual intra-modal conditions were not significantly different (all ps>0.1). These results rule out the possibility that the observed segregations in neural representations of auditory and visual stimuli (**Figure 3A-E**) were due to their differences in other properties than their subjective values.

Secondly, similar to the analysis of the absolute differential value, we analysed the mean reaction time (RT) data for the two types of feedback in the value and the control tasks. A two-way repeated-measures ANOVA of RTs with task and feedback as factors revealed no significant main or interaction effects (p-values > 0.05). However, a trend was found for the main effect of task F[2,38] = 4.76, p = 0.06, reflecting faster responses in the control compared to the value task. Overall, the mean RT in the control task (787.5(±0.0613) ms), where participants had to simply follow instructions for decision-making was shorter than the mean RT in value task (824.4(±0.0635) ms).

Intuitively, a systematic decrease in the mean RTs of value task along with an increase in the absolute differential values (from no-reward to reward feedbacks), would indicate that participants take more time to reach a decision during difficult choice trials (when both options were perceived as having approximately equal values) in comparison to easy choice trials (when one option was clearly more valuable than the other). In the value task, mean (±s.e.m.) RTs decreased from 828.3(±0.0634) ms (no reward/blue feedbacks) to 819.7(±0.0638) ms (reward/yellow feedbacks). On the contrary, in the control task, mean RTs increased from 786.7(±0.0620) ms (keep/blue feedbacks) to 792.9(±0.0580) ms (switch/yellow feedbacks). Although insignificant, the latter effect implies an obvious fact that participants took less time when they had to keep their past choice in comparison to making a switch. Neither the main effect of feedback type on RTs nor their interaction with the task however reached significance, based on ANOVA (Fs<1, p>0.1).

### Comparison of the value and the control task

In addition to the value task, we also inspected the control task using the same contrasts that were used to detect modality-specific and modality-general representations shown in **Figure 3**. Interestingly, we found that in the control task there were activations (**Figure S4** and **Table S3**) in vmPFC that overlapped with modality-general representations that were found in the value task. This observation indicates that a task with comparable choice structure, but no valuation requirement also involves vmPFC, underscoring the role of this region as a general comparison and choice computation region. No activations were found at the OFC with these contrasts at the reported threshold.

### Effective Connectivity Analysis

The model space shown in **Figure 4** and **Figure 5** was developed over a base model comprising driving inputs and intrinsic connections, which did not vary with the experimental conditions. The rest of the models differed from each other over modulatory connections, which depended on the experimental conditions **(Figure 4A)**. In the base model, intrinsic connections were defined between every pair of nodes in the network and as self-connections. In **Table S4** and **S5**, we show the estimated strength of the intrinsic connectivity (**Table S4**) and driving inputs (**Table S5**) of the winning model.

---

**Figure S1.**
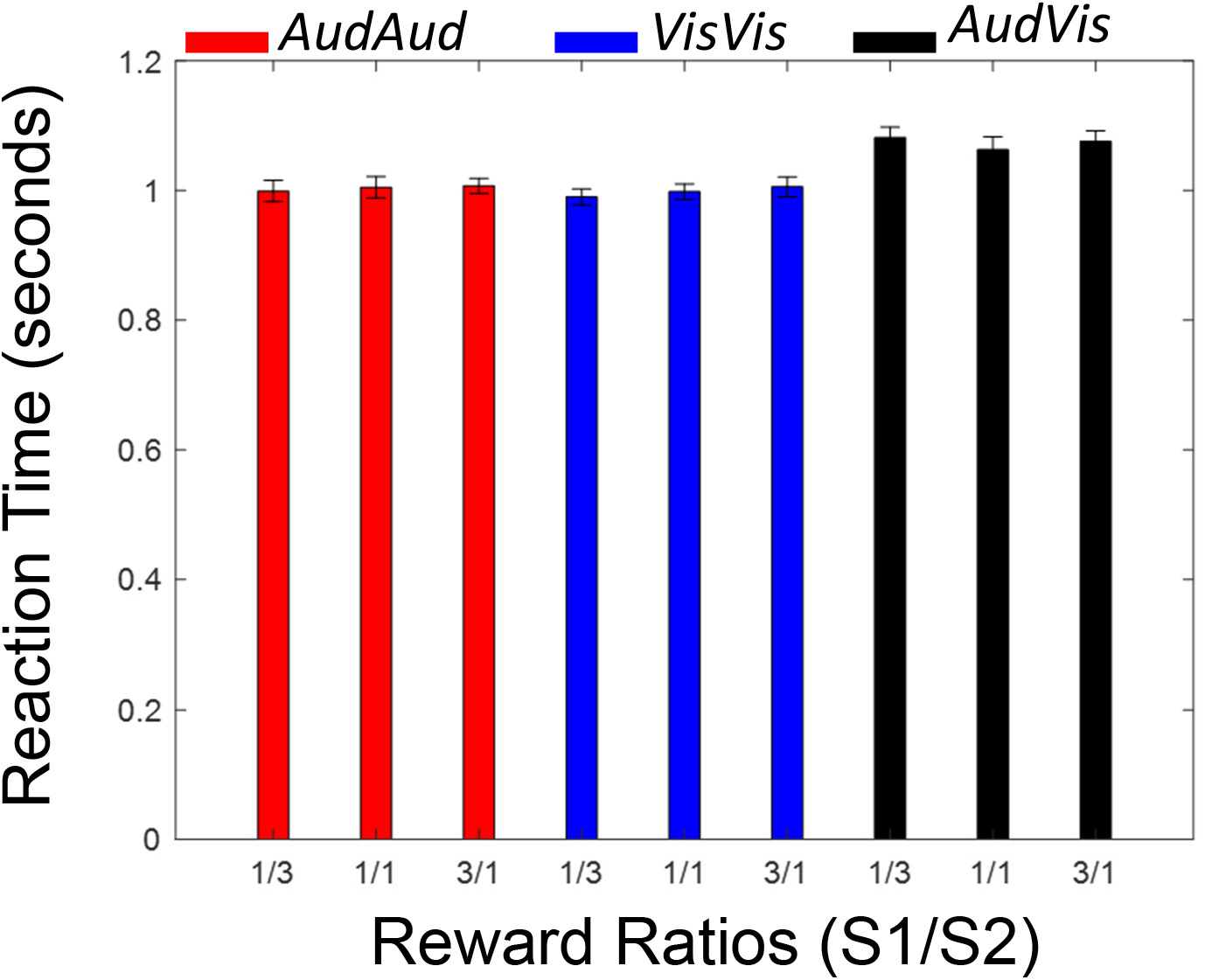
Reaction Times in the value task. Mean RTs across participants for each reward ratio {1:3, 1:1, 3:1} of options *S*_1_: *S*_2_ separately for each modality condition of the value task (*AudAud*, *VisVis* and *AudVis*).

**Figure S2.**
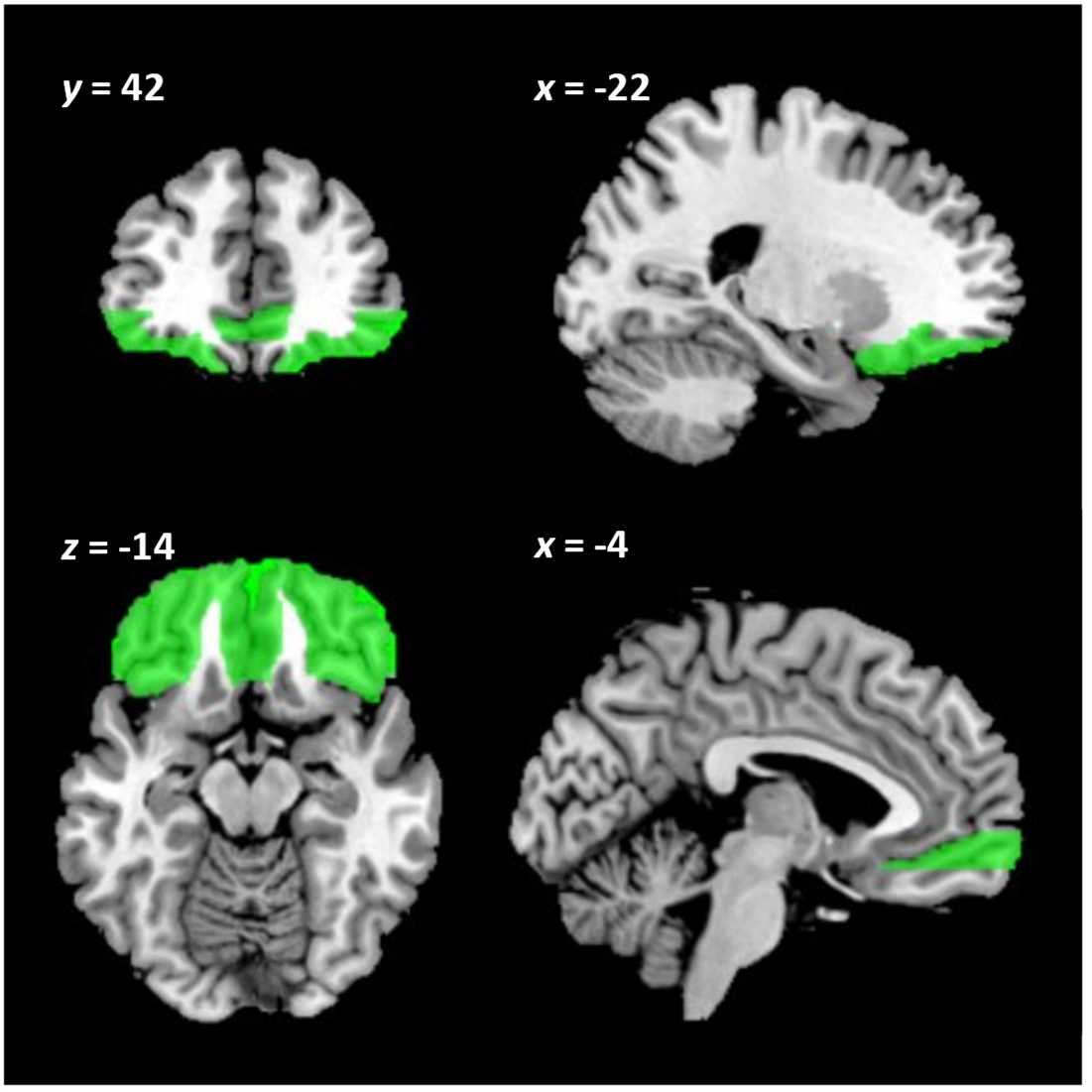
Anatomical definition of frontal valuation areas. The search volume used for multiple comparisons correction consisted of anatomical parcellations of the orbital surface of frontal gyrus as defined in automated anatomical labelling (AAL) atlas (Rolls et al., 2020, 2015). The search volume, comprised the anatomical parcellations of orbital surface in the following format, ROI name (abbreviation): Superior frontal gyrus - medial orbital (PFCventmed); Medial orbital gyrus (OFCmed); Anterior orbital gyrus (OFCant); Posterior orbital gyrus (OFCpost); Lateral orbital gyrus (OFClat), for the detailed description of these areas see Table 2 in Rolls et al. 2015, 2020 (Rolls et al., 2020, 2015).

**Figure S3.**
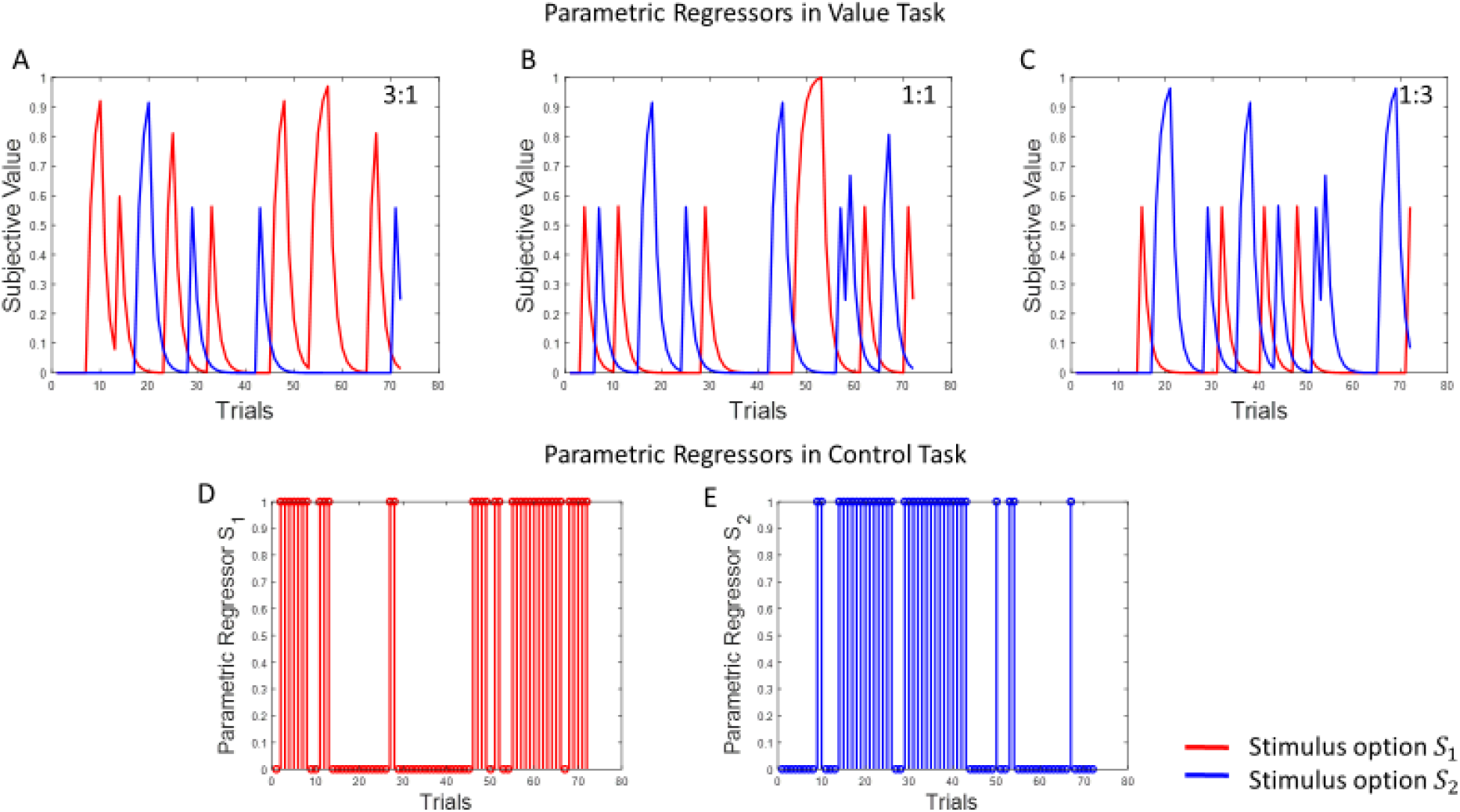
Illustration of the time course of parametric regressors in the value and control tasks. Parametric regressors used for the fMRI analysis are shown for a single participant. (**A-C**) For the value task, subjective values (SVs) of each option (*S*_1_, *S*_2_) are shown across trials in a block with reward ratios of 3:1, 1:1, 1:3, respectively. SVs were calculated based on the computational modeling of the behavioural data (see Material and Methods in the main text). (**D-E**) For the control task, where instructions were passively followed across trials in a block, a weight of 0 or 1 was assigned to each option. The weights assigned to *S*_1_and *S*_2_ in the control task were determined based on the schema shown in **Table S1**.

**Figure S4.**
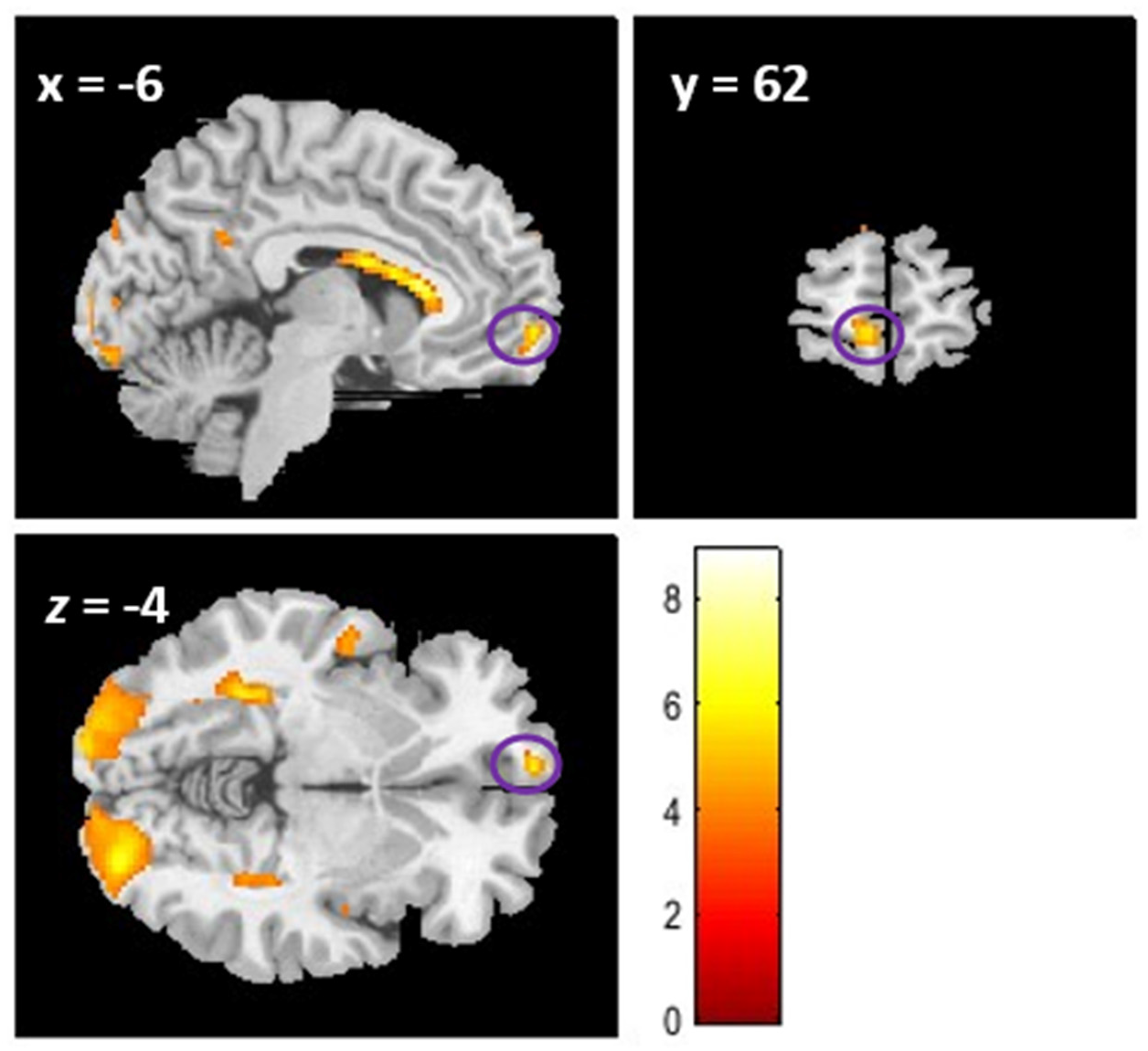
Activations in vmPFC in the control task. The activations correspond to the effect of parametric regressors across all modality conditions. The results are shown at the whole-brain uncorrected level of P < 0.001.

**Figure S5.**
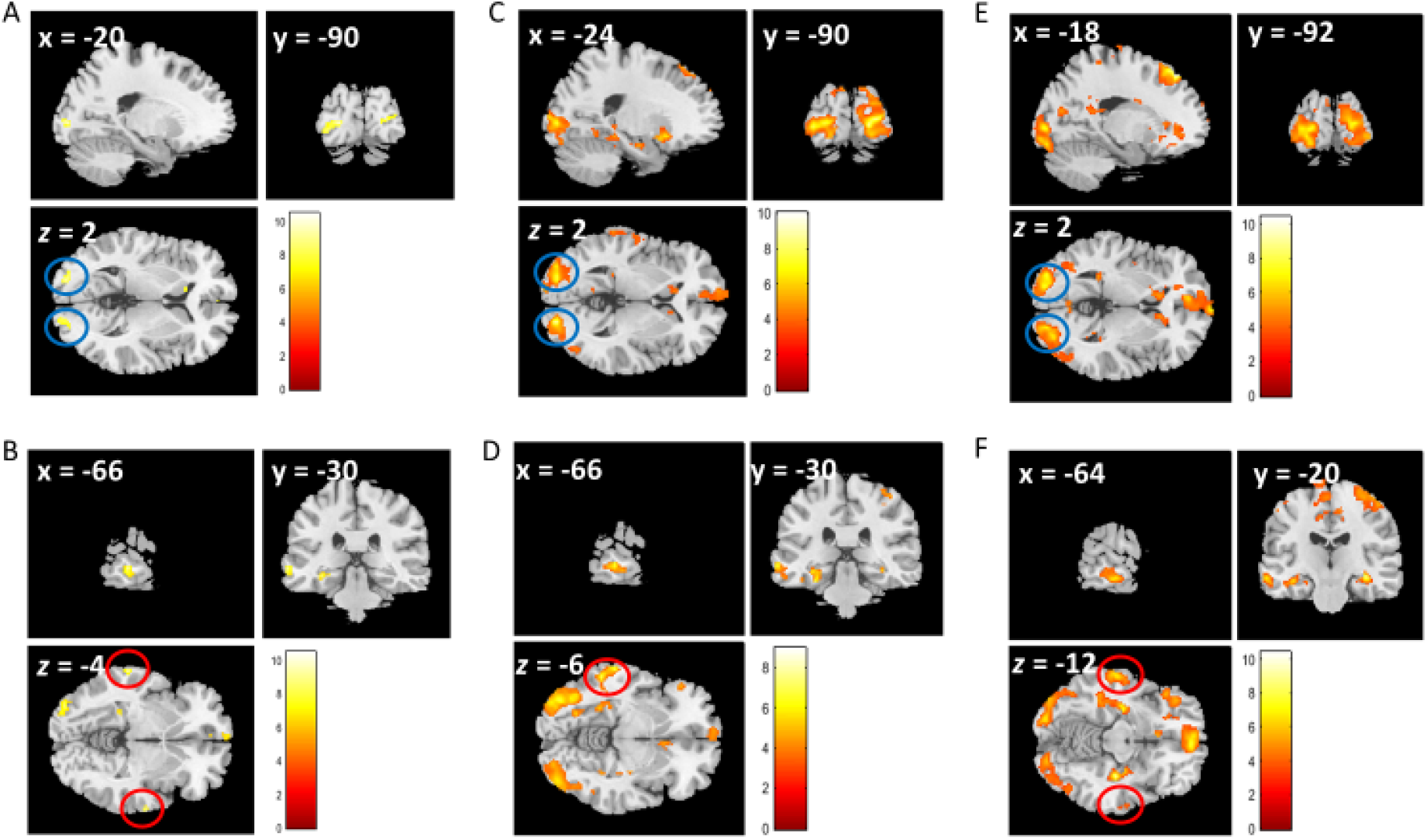
Stimulus value representations in sensory cortices: (**A**) Visual sensory cortex. (**B**) Auditory sensory cortex. All cluster activations shown in (A) and (B) were significant at whole-brain FWE corrected P < 0.05 and are estimated across all conditions (*AudAud, VisVis and AudVis*). (**C**) When the value modulations were inspected for individual conditions separately (whole-brain uncorrected level of P < 0.001), examination of the value parametric regressors in visual modality (i.e., contrast *intravisSV* > 0) revealed activations in the visual sensory cortex (cluster peaks at (-24, -90, 2) and (18, -94, 8)) but no activation in the auditory sensory cortex were found. (**D**) Similarly, for the contrast *intraaudSV* > 0, we found activations in the auditory sensory cortex (cluster peak at (-66, -30, -6)) but no activation in the visual sensory cortex. This result indicates that each sensory cortex was maximally activated when the value of a stimulus from its specific modality was processed. However, for the latter contrast (*intraaudSV* > 0), we also found activations in higher visual areas (occipitotemporal cortex) with cluster peaks at (-36, -66, -8) and (42, -76, -12) that were distinct from activations found for the contrast *intravisSV* > 0 which were in early visual areas in the occipital cortex (anatomical definitions are based on https://neurosynth.org/). (**E-F**) For the contrast *interaudvisSV* > 0, we found activations in both visual (cluster peaks at (-18, -92, 2) and (22, -90, 6)) and auditory (cluster peaks at (-64, -20, -12) and (66, -10, -4)) cortex, as in audio-visual condition trial-by-trial subjective values are updated individually for both auditory and visual options (whole-brain uncorrected level of P < 0.001). In all figures, crosshairs are placed at the left hemisphere cluster peak. The blue and red circles mark activations in the visual and auditory cortex, respectively.

**Table S1.**
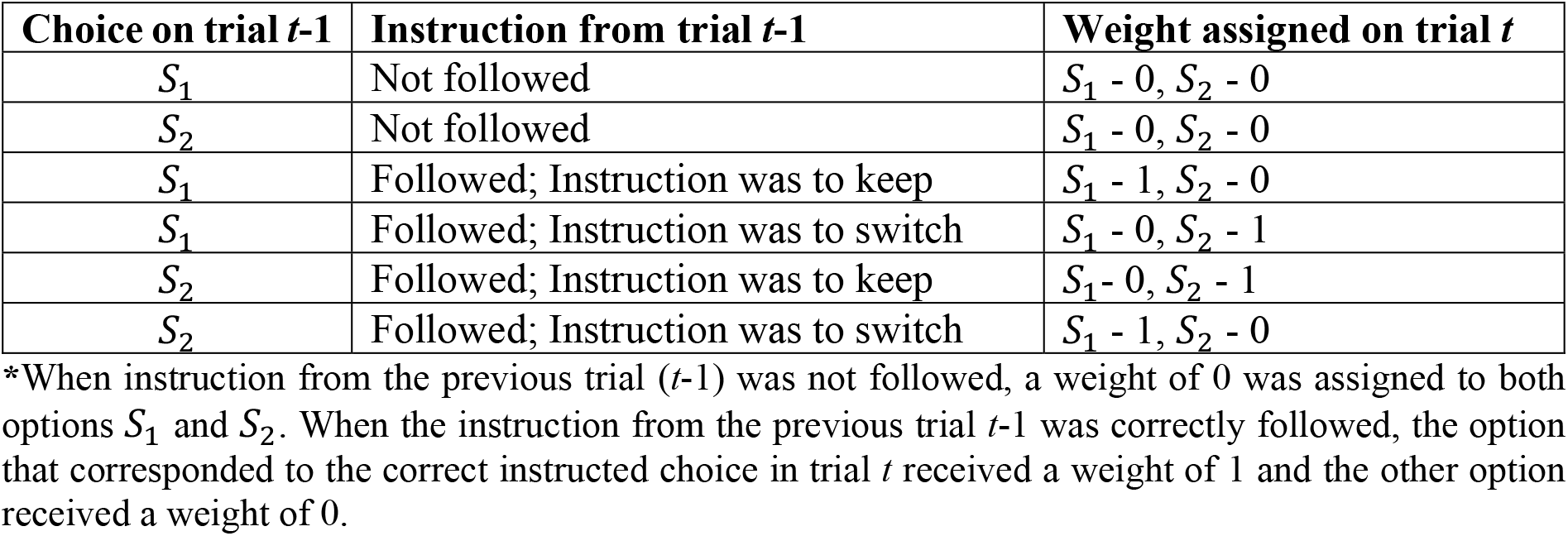
Definition of parametric regressors for the control task*.

| **Table S1. Definition of parametric regressors for the control task\*** |  |  |
| --- | --- | --- |
| **Choice on trial $t-1$** | **Instruction from trial $t-1$** | **Weight assigned on trial $t$** |
| $S_1$ | Not followed | $S_1 - 0, S_2 - 0$ |
| $S_2$ | Not followed | $S_1 - 0, S_2 - 0$ |
| $S_1$ | Followed; Instruction was to keep | $S_1 - 1, S_2 - 0$ |
| $S_1$ | Followed; Instruction was to switch | $S_1 - 0, S_2 - 1$ |
| $S_2$ | Followed; Instruction was to keep | $S_1 - 0, S_2 - 1$ |
| $S_2$ | Followed; Instruction was to switch | $S_1 - 1, S_2 - 0$ |

\*When instruction from the previous trial ( $t-1$ ) was not followed, a weight of 0 was assigned to both options $S_1$ and $S_2$ . When the instruction from the previous trial $t-1$ was correctly followed, the option that corresponded to the correct instructed choice in trial $t$ received a weight of 1 and the other option received a weight of 0.

**Table S2.** Results of the post-hoc pairwise comparisons of choice ratios between different modalities.

|  |  |  |  |  |
| --- | --- | --- | --- | --- |
| Reward Ratio | Modality-1 | Modality-2 | Difference | pValue |
| 1:1 | Auditory | Visual | 0.0417 | 0.9857 |
| 1:1 | Auditory | AudioVisual | 0.1069 | 0.0626 |
| 1:1 | Visual | Auditory | -0.0417 | 0.9857 |
| 1:1 | Visual | AudioVisual | 0.0651 | 0.3084 |
| 1:3 | Auditory | Visual | 0.0548 | 0.3278 |
| 1:3 | Auditory | AudioVisual | 0.0701 | 0.2903 |
| 1:3 | Visual | Auditory | -0.0548 | 0.3278 |
| 1:3 | Visual | AudioVisual | 0.0154 | 1.0 |
| 3:1 | Auditory | Visual | 0.0325 | 1.0 |
| 3:1 | Auditory | AudioVisual | 0.1424 | 0.0873 |
| 3:1 | Visual | Auditory | -0.0325 | 1.0 |
| 3:1 | Visual | AudioVisual | 0.1098 | 0.0156* |

\* Indicates significance at $p < 0.05$

**Table S3.** Univariate contrasts inspected in the control task.

| Contrast | Region | *x* | *y* | *z* | *t*(19) | *k* | *SVFWE corr P* |
| --- | --- | --- | --- | --- | --- | --- | --- |
| *Control_intraAud* > 0 | vmPFC | -8 | 62 | -4 | 4.49 | 11 | 0.300 |
| *Control_intraVis* > 0 | vmPFC | -6 | 64 | -6 | 4.37 | 12 | 0.287 |
| *Control_interAudVis* > 0 | vmPFC | -4 | 56 | -12 | 2.87 | 8 | 0.966 |

MNI coordinates (*x*, *y*, *z*) and *T* value corresponds to the local maxima peak of the cluster activations (cluster labels are from AAL atlas). Statistical maps were assessed for cluster-wise significance using a cluster-defining threshold of $t(19) = 3.58$ , $P = 0.001$ ; and using small volume correction within the frontal search volume but cluster-level SVFWE corrected *P*-values were above threshold *P*-value 0.005 (as shown in last column of the table).

**Table S4.**
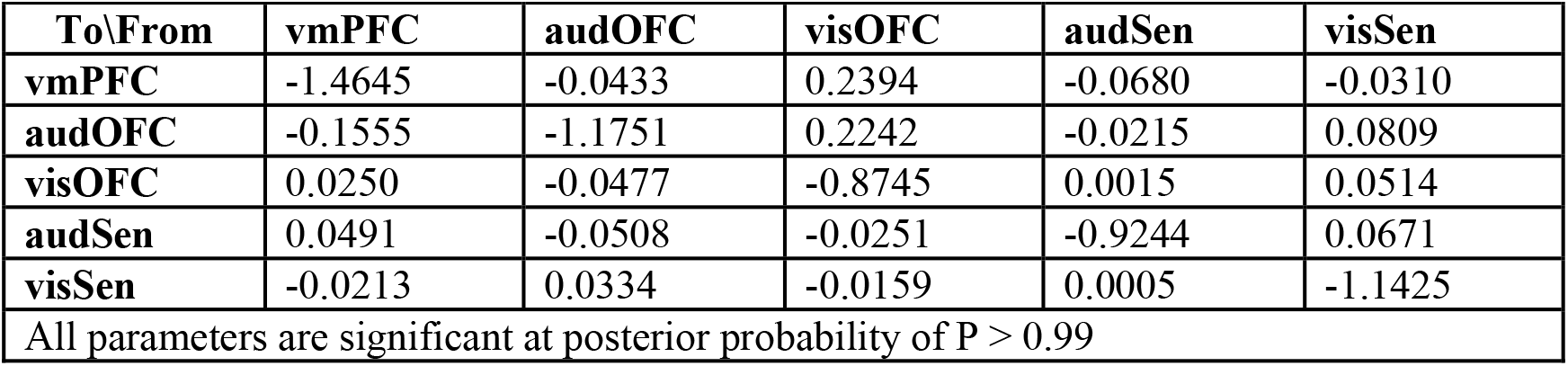
Intrinsic connectivity parameters in the network including self-connectivity.

|  |  |  |  |  |  |
| --- | --- | --- | --- | --- | --- |
| To\From | vmPFC | audOFC | visOFC | audSen | visSen |
| vmPFC | -1.4645 | -0.0433 | 0.2394 | -0.0680 | -0.0310 |
| audOFC | -0.1555 | -1.1751 | 0.2242 | -0.0215 | 0.0809 |
| visOFC | 0.0250 | -0.0477 | -0.8745 | 0.0015 | 0.0514 |
| audSen | 0.0491 | -0.0508 | -0.0251 | -0.9244 | 0.0671 |
| visSen | -0.0213 | 0.0334 | -0.0159 | 0.0005 | -1.1425 |
| All parameters are significant at posterior probability of $P > 0.99$ | | | | | |

**Table S5.** Driving input influence parameters on sensory ROIs of the network.

|  |  |  |  |  |
| --- | --- | --- | --- | --- |
| ROIs\Driving Input* | intraAud | intraVis | interAud | interVis |
| audSen | 0.0042 | 0 | 0.0226 | 0.0224 |
| visSen | 0 | -0.0025 | -0.0035 | 0.0240 |
| All parameters are significant at posterior probability of $P > 0.99$ | | | | |
| *For trial-types of different conditions of the value task as described in the methods section on EC |  |  |  |  |

## Notes

### Competing Interest Statement

The authors have declared no competing interest.

### Summary of Updates

1) Introduction has been streamlined. 2) Methods has been extensively revised to increase the clarity. 3) Figure 3 is updated: showing the generalization of modality-specific representations across conditions. 4) Figure 4 is updated: to check the effective connectivity explicitly for the existence of cross-modal connections.

